# Investigating spatial dynamics in spatial omics data with StarTrail

**DOI:** 10.1101/2024.05.08.593025

**Authors:** Jiawen Chen, Caiwei Xiong, Quan Sun, Geoffery W. Wang, Gaorav P. Gupta, Aritra Halder, Yun Li, Didong Li

## Abstract

Spatial omics technologies revolutionize our view of biological processes within tissues. However, existing methods fail to capture localized, sharp changes characteristic of critical events (e.g. tumor development). Here, we present StarTrail, a novel gradient based method that powerfully defines rapidly changing regions and detects “cliff genes”, genes exhibiting drastic expression changes at highly localized or disjoint boundaries. StarTrail, the first to leverage spatial gradients for spatial omics data, also quantifies *directional* dynamics. Across multiple datasets, StarTrail accurately delineates boundaries (e.g., brain layers, tumor-immune boundaries), and detects cliff genes that may regulate molecular crosstalk at these biologically relevant boundaries but are missed by existing methods. For instance, StarTrail precisely pinpointed the cancer-immune interface in a HER2+ breast cancer dataset, unveiled key cliff genes including a potential prognostic biomarker *IGSF3*, highlighting NK-, B-cell mediated immunity, and B cell receptor signaling pathways missed by all spatial variable gene methods attempted. StarTrail, filling important gaps in current literature, enables deeper insights into tissue spatial architecture.

## 1 Main

Spatial data, with its inherent geographical, topological or geometric coordinates, enables structured views of cellular and inter-cellular dynamics and allows researchers to discern patterns, relationships, and changes over different spatial scales. Leveraging spatial information has proven invaluable in numerous fields including biomedical imaging studies, environmental science and computational biology [see, for e.g., 1–3]. In recent years, spatial omics technologies have emerged as valuable tools in biological and biomedical studies [4]. Their significance lie in the unique capability to provide omics measurements while simultaneously retaining spatial location information, e.g. gene expression in spatial transcriptomics (ST), and protein expression in spatial proteomics. Notable technologies in this domain include 10X Visium [5], Slide-seqV2 [6], and MERFISH [7] for ST, and CODEX [8] earmarked for spatial proteomics. The dual insight has transformed our understanding of cellular organization within tissues.

The ST literature has seen several focused developments that have garnered attention and gained popularity. Methodology employed in the detection of spatially variable genes (SVGs) is one of the most popular [see, e.g., 9, 10] areas. It aims to detect genes that exhibit differential expression (DE) across spatial locations. SVG detection methods enable the identification of important genes that are associated with tissue architecture, function, or pathology, based on their spatial expression patterns. Note that SVGs and DE genes (DEGs) have been used exchangeably in ST literature. Here we use SVGs for genes exhibiting spatial changes without the need to pre-define spatial regions and use DEGs for genes showing differences between two pre-defined regions. Besides SVG and DEG detection, there is an increasing emphasis on developing methods to detect spatial clusters, also often referred to as spatial domains [see, e.g., 11–14], specifically using histology images and spatial coordinates. These spatial clustering methods aim to reliably represent the natural biological patterns found within tissues.

However, at least three notable limitations are present in the aforementioned methodologies. First, although methods for identifying SVGs can reveal genes that vary across space, they are unable to pinpoint the exact locations of these variations. In the clustering context, the spatial domains defined can vary substantially depending on which genes are selected for analysis. Second, no existing method is sufficiently powerful to identify omics features (e.g., genes, proteins) that exhibit drastic changes along highly localized or disjoint boundaries. We name such genes cliff genes, distinct from SVGs, which typically manifest global changes or at least across larger areas. Third, existing methods provides no directionality information. In other words, no method quantifies how exactly the expression of any given gene changes in the 2-dimensional (2D) space. Owing to these methodological gaps, the underlying intricacies that drive the manifestation of localized boundaries in spatial omics data remain largely unexplored. Many current approaches rely on DE analyses at specific boundary intersections [15, 16], but this barely touches on the depth of spatial dynamics involved.

To address these limitations, we introduce StarTrail (Scalable spaTiAl gRadienT pRocess Approx-Imation utiLizing nearest-neighbor Gaussian process). StarTrail is the first method that leverages 2D spatial gradient for the analysis of spatial omics data. Spatial gradients, a concept borrowed from calculus, measuring rate of changes across spatial locations, offer perspectives complementary to raw omics measurements and thus into nuances of spatial data changes missed by existing approaches. Specifically in response to the three aforementioned limitations, StarTrail provides quantitative, directional, super-resolution gradient estimates that illuminate more precisely where and how changes occur. It is important to note that our use of the term ‘gradient’ specifically refers to its mathematical sense as the derivative of a function, describing the rate of changes in an omics feature relative to spatial coordinates. This definition is distinct from the more philosophical or intuitive use of ‘gradient’ found in some other studies [see, e.g., 17, 18]. Our focus is on the concrete and quantitative aspects of spatial data variation.

As a motivating example, in cancer samples, the tumor region(s) can be rather small (e.g., newly developed or developing cancerous regions) and disjoint (multiple primary or a combination of primary and metastatic regions) with each having its own localized boundary. Existing methods proposed to identify SVGs are under-powered to identify omics features that manifest drastic changes along these boundaries because they can easily be an negligible portion of the entire tissue space examined. These boundaries, however, frequently coincide with vital biological junctures, such as the interface that distinguishes diseased from healthy tissues or areas of distinct biological functions [see, e.g., 17]. Complemented with histology images, gradient analyses are further enriched, e.g.,g enabling the analysis of gradient along pathologist-annotated boundaries. Previous research primarily concentrated on constructing slide topological maps using low-dimensional features, typically one-dimensional (1D) attributes such as pseudo-time or iso-depth [see, e.g., 17, 19–22]. These studies often leaned on stringent assumptions, such as assuming that every gene’s gradient direction is either null or aligned with the 1D latent variable [19], or the gradient calculation depends heavily on the choice of the model [20]. However, accurately inferring gene-wise spatial gradients [19] and appropriately applying gradients to understand spatial dynamics of each gene expression remain as open and challenging tasks.

StarTrail is highly flexible and versatile: it can accommodate various spatial omics features, encompassing genes, proteins, as well as annotations or derived features such as estimated cell type proportions. StarTrail harnesses spatial gradients to project the vector field of each omics feature, pinpointing regions of the most rapid change by their heightened gradient values. This enables the delineation of omics boundaries through principal curve [23] (see section 2.1 below). Moreover, when boundaries are pre-defined (e.g., based on pathologists’ annotations), StarTrail adeptly calculates gradients along their normal vectors (wombling, see section 2.1 below), defining “cliff genes” whose gene expressions shift most sharply across the boundaries. Incorporating gradient metrics and Wombling analysis [24], Star-Trail also enhances downstream analysis potentials. For instance, StarTrail can categorize SVGs based on their gradients relative to predefined boundaries. Additionally, StarTrail can bolster clustering analysis by leveraging the gradient metric, providing more refined insight into spatial data structures, compared with only utilizing the original space. Notably, StarTrail achieves a marked increase in computational efficiency, compared to the original spatial gradient process [25, 26], facilitating rapid and sophisticated spatial data analysis.

We demonstrate the versatility of StarTrail on a spectrum of spatial omics data types and datasets, including omics data generated by technologies such as 10X Visium [5], Spatial Transcriptomics [5], and CODEX [8], as well as derived features such as estimated cell type proportions.

## 2 Results

### 2.1 Scalable spatial gradient process via StarTrail

The overview of StarTrail is illustrated in Fig. 1. To understand the relationships between spatial co-ordinates *s* and omics feature *Y*, StarTrail harnesses the Gaussian Process (GP), *Z*. Designed to be versatile, StarTrail operates seamlessly with both 2D and 3D coordinates. The input data, *Y* can encompass various spatial omics or spatial annotations, including metrics such as gene expression, protein abundance, or cell type proportions. In this illustration, we use gene expression as an example. The subsequent gradient is adeptly modeled using a spatial gradient process denoted by ∇*Z*. This forms a joint distribution of [*Z*, ∇*Z*], with cross-covariance characterized by *K* and its derivatives up to the second order [see, for e.g., 25–27]. While GPs offer the advantageous property wherein their derivative remains a GP, the large sample size (e.g. the number of spots/cells in ST) in spatial omics data makes the use of the original spatial gradient process untenable [25, 26]. For instance, it takes over 40 hours to fit the original spatial gradient process on ∼ 3000 spots for one gene. To address this, StarTrail introduces a scalable spatial gradient process which utilizes finite differences to approximate gradients. Inspired by both plug-in GP [28] and the nearest-neighbor (NN) GP [29, 30], StarTrail establishes a NNGP model. The integration of NN facilitates a significant computational economy, reducing the cost of fitting the GP from O(*n*^3^) to O(*nm*^3^), where *n* represents the total sample size and *m* is the number of neighboring locations used [29, 30].

**Figure 1.**
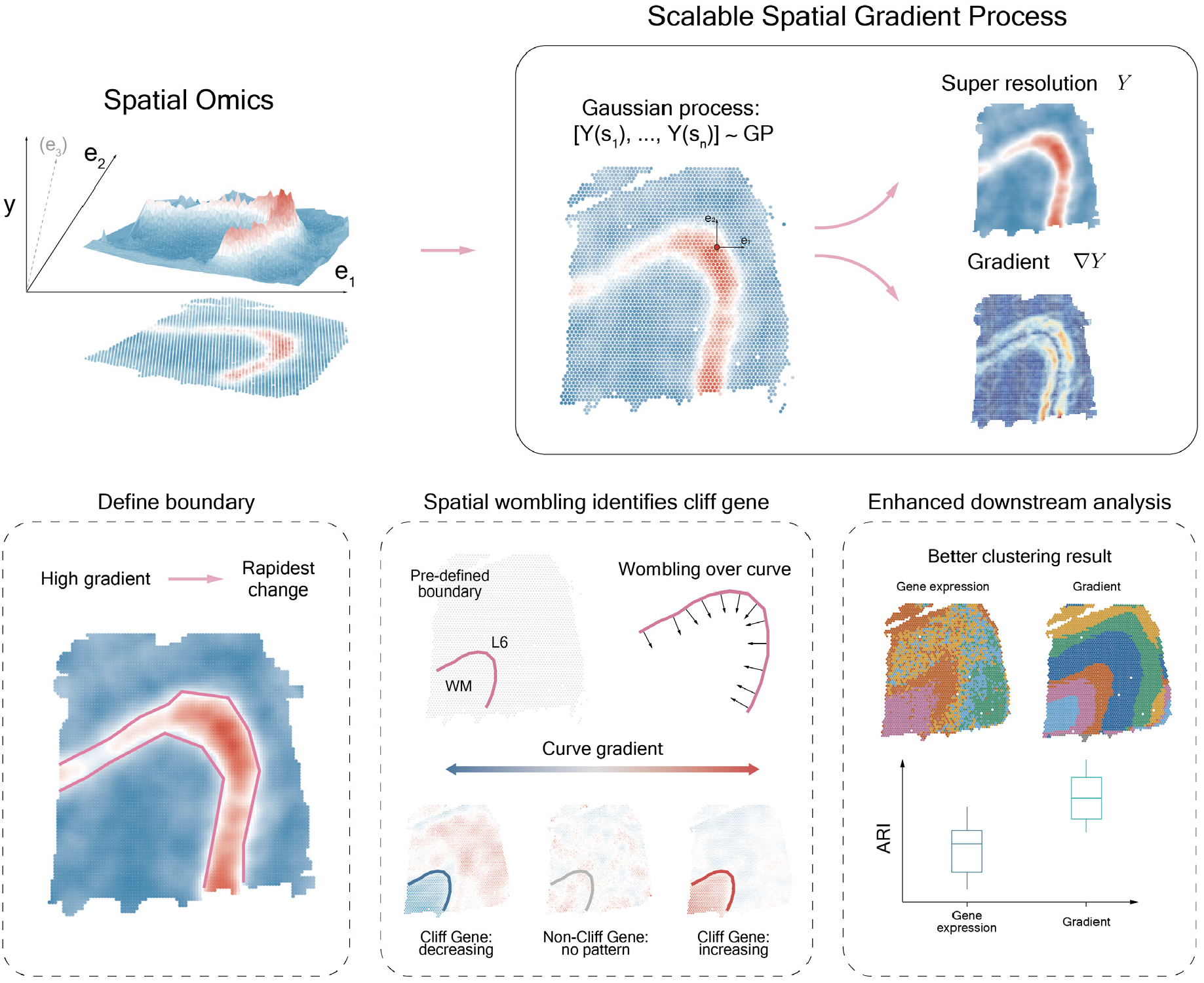
StarTrail overview. StarTrail takes spatial omics data as its input (top left). Utilizing a scalable spatial gradient process, StarTrail models the intricate relationship between spatial coordinates *s* and omics features *Y*, efficiently inferring gradients across a tissue region (top right). With the inferred gradients, StarTrail can identify boundaries by pinpointing areas exhibiting the highest gradient values (bottom left). Additionally, StarTrail employs Wombling analysis across predefined boundaries to detect “cliff genes”—genes with significant expression changes at these boundaries (bottom middle). Moreover, these gradients could be leveraged to enhance downstream analyses, such as clustering (bottom right).

Leveraging the advantages of GPs, StarTrail offers the ability to infer spatial omics data in previously unmeasured regions, thereby enhancing the resolution to any desired level. Utilizing the inferred gradient, ∇*Z*, StarTrail enables a detailed examination of the spatial dynamics. More precisely, it enables the evaluation of gradients in any chosen direction. For preliminary analysis, for instance, the process initiates by deducing the gradient along the directions represented by *e*_1_ := (1, 0) (horizontal direction) and *e*_2_ := (0, 1) (vertical direction), respectively. The gradient at every point is then calculated as 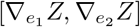, from which we define the gradient flow map which characterizes the direction of local change for each spot/cell. Following this, the *L*^2^-norm of gradients 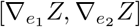 serves to determine the magnitude of the gradient. This nuanced approach enables us to categorize the dynamics of omics data—rapid changes are indicated by high gradient values, while areas exhibiting slow changes are characterized by low-magnitude gradients. For omics features exhibiting swift alterations across a region, boundaries can be identified along the area of maximal gradient, by using a non-parametric curve fitting algorithm to delineate principal curves [for more details, see 23].

In addition to the ability to infer gradients and delineate boundaries, we recognize that it is equally crucial to evaluate the directional gradients along a given boundary. The boundary is usually annotated by pathologists or experts with domain knowledge. Delving into omics alterations across specific boundaries provides invaluable biomedical insights, particularly in scenarios where these boundaries demarcate pivotal biological structures or lurking conditions. To leverage pathologist-annotated boundaries or boundaries informed by external information, such as cell type proportion demarcations, we engage the “Wombling” technique. This approach involves statistical inference on directional derivatives evaluated along the normal direction (also known as the orthogonal direction) of pre-designated boundary segments. Subsequent integration across the entire curve produces a valid measure that can be used to track whether a curve forms a “wombling boundary” [see 24, Section 3.3, for more detail]. Upon analyzing the curve gradient inferred using wombling, we introduce the concept of a “cliff gene”. A cliff gene designates a gene that exhibits pronounced shifts across specific boundaries. Analogously, we can expand this concept to other domains and similarly define a “cliff cell type” or a “cliff protein”, each signifying an omics feature that manifests significant variations over demarcated regions.

Utilizing the inferred gradients provides perspectives on spatial dynamics beyond the original feature space, enhancing and enriching downstream analyses. Using StarTrail inferred spatial metrics, we can precisely link SVGs to specific spatial boundaries based on their gradients. This granularity enables a more targeted and precise analysis. Furthermore, the integration of gradient metrics enhances clustering analysis, delineating distinct clusters based on spatial intricacies.

### 2.2 StarTrail accurately estimates spatial gradient and defines rapid change area

We first benchmark the performance of StarTrail using a series of simulation tests designed to mirror a variety of spatial patterns. They range from trigonometric functions (Extended Data Fig. 1, first row), to common patterns including linear gradients and hot-spots, as well as regional patterns (Extended Data Fig. 1, second row-last row). For each simulated scenario, we calculated gradients along the directions *e*_1_ = (1, 0) and *e*_2_ = (0, 1). The results show that StarTrail achieves high accuracy in recovering the gradients along these axes. To pinpoint areas of rapid changes in any given direction, we computed the *L*^2^-norm of the gradients (referred to *L*^2^ in the following context). This metric successfully identified regions with the highest gradient magnitude, corresponding to the points of most significant changes in our simulation models. Linear gradients and hot-spots represent patterns of changes gradually from one constant to another constant, while regional patterns exhibit abrupt jumps from one constant to another. Consequently, we have chosen to use *L*^2^ as the indicator to mark areas undergoing rapid changes when applying StarTrail to real-world data analysis.

After confirming StarTrail’s efficacy through simulations, we turned our attention to real-world spatial omics datasets, starting with ST data. We first analyzed 10x Visium data from human dorsolateral prefrontal cortex (DLPFC)[16]. The DLPFC dataset is known for its well-defined seven-layer architecture, including layers L1 through L6 and white matter (WM), making it an exemplary instance for studying layer-enriched gene expression. The spatially resolved expression has important clinical implications for neurological disorders such as autism spectrum disorder and schizophrenia [16]. StarTrail aptly captures the intricate relationship between spatial coordinates and gene expression levels, as well as their spatial gradients. This capability is demonstrated through the precise modeling of several key layer marker genes (Fig. 2b-d, other example genes in Extended Data Fig. 2). *MBP*, a marker gene enriched in WM, shows spatial gradients with the highest *L*^2^ values concentrated at the boundary between WM and layer L6. StarTrail further identifies the boundary using the principal curve fitted to the points with *L*^2^ exceeding its 90% quantile. The resulting boundary (Fig. 2b) accurately defines the zone of the most dramatic expression changes. Additionally, the direction of these gradients is effectively summarized by aggregating the horizontal and vertical gradients ([31], Fig. 2c). The gradient flow map generated from this approach provides an insightful visualization of gene expression dynamics, visualizing the change direction of gene expression, and pinpointing locations of extreme expression (singularities), where neighboring gradients converge (indicating maximal expression) or diverge (indicating minimal expression). StarTrail also accurately estimated spatial gradients for other layer-specific markers such as *PCP4* for layer L5 and *HPCAL1* for layer L1, further demonstrating the robustness and accuracy of StarTrail in deciphering spatial expression patterns. Moreover, one fundamental advantage of employing a spatial gradient process lies in its ability to construct a spatial model between spatial locations and their corresponding omics feature as well as gradients. Consequently, StarTrail is equipped to extrapolate spatial omics features to areas that were not originally sampled, effectively filling in the gaps and providing a more complete super-resolutional picture of the spatial dynamics (Fig. 2b).

**Figure 2.**
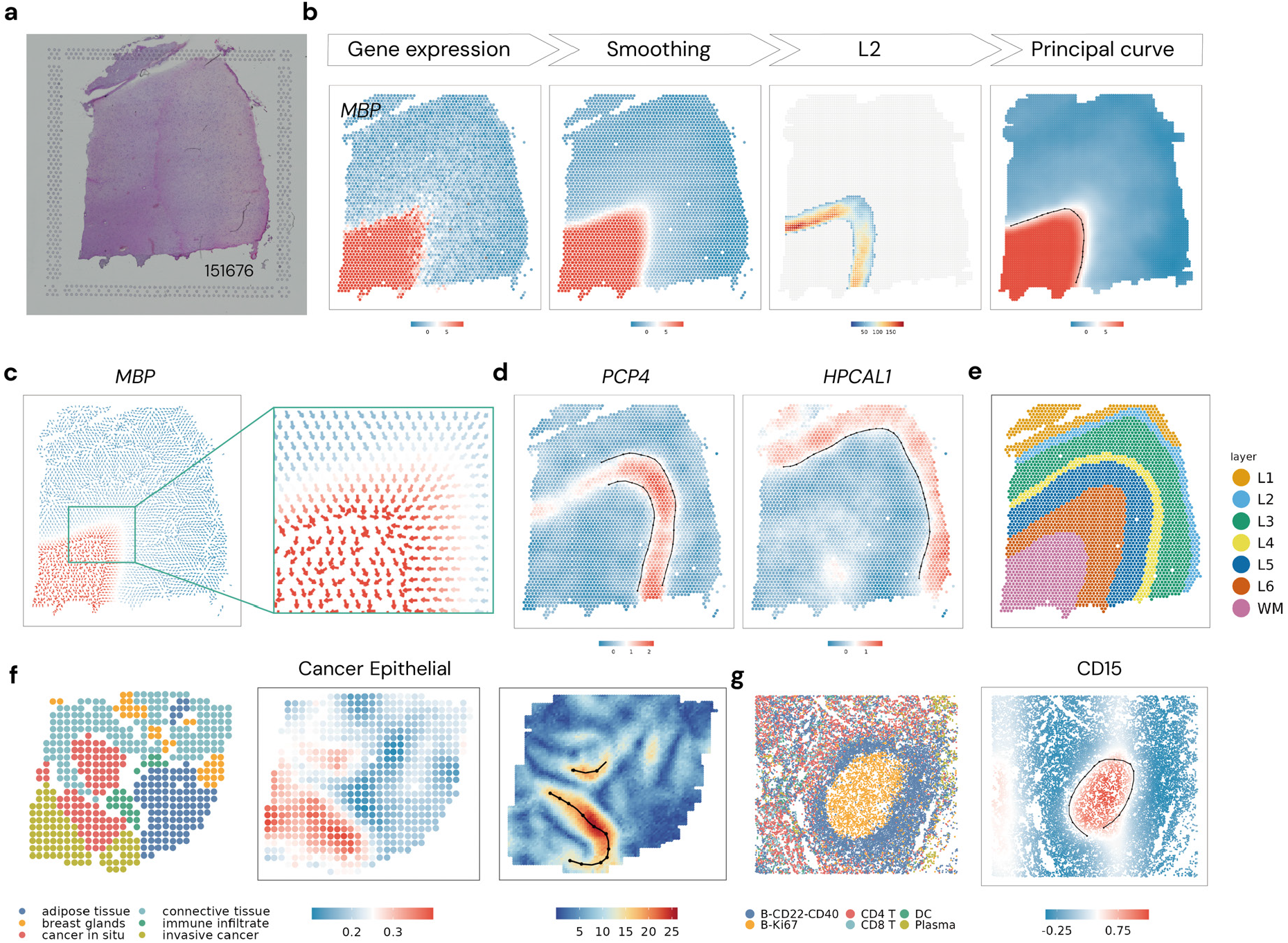
Gradient analysis. **DLPFC**: (a) Histology image of DLPFC slide 151676. (b) A workflow outlines the process of detecting gene expression boundaries, using *MBP* as an example. StarTrail first smoothes raw gene expression and estimates the spatial gradients, and then fits a principal curve to spots where the *L*^2^ norm of the gradients exceeds the 0.9 quantile, effectively delineating the gene expression boundary. (c) Gradient flow map for *MBP*, arrows colored by smoothed and normalized expression (d) Inferred boundaries for *PCP4* and *HPCAL1*, points colored by smoothed and normalized expression (e) Pathologist annotation. **HER2+ breast cancer**: (f) left: pathologist annotation, middle: estimated cancer epithelial cell proportion, right: estimated *L*^2^. **CODEX**: (g) left: cell type assignment, right: detected boundary for CD15, points colored by smoothed and normalized abundance.

Moving to a finer-resolution and non-layered dataset, we next applied StarTrail on HER2+ breast cancer data from the Spatial Transcriptomics platform (focusing on slide H1 shown in Fig. 2f). Breast cancer encompasses a variety of subtypes, among which HER2-positive (HER2+) subtype is particularly known for the overexpression of *ERBB2*, commonly referred to as *HER2* [32]. The dataset was annotated by pathologist into six distinct histo-pathological regions: noninvasive ductal carcinoma in-situ (DCIS), invasive breast cancer (IBC), adipose tissue, immune infiltrate, breast glands, or connective tissue (Fig. 2f left panel). As mentioned, a key advantage of StarTrail is its adaptability to various spatial measurements. For this dataset, we applied StarTrail to model the spatial distribution of cell type proportions inferred by RCTD (Fig. 2f middle panel) [33]. Cancer epithelial cells, as expected, are predominantly located within cancerous regions [32, 34, 35]. StarTrail’s inference of spatial gradients showed accurate delineation, particularly with high *L*^2^ values around the DCIS zones (Fig. 2f right panel). Notably, the steepest gradients were identified at the interface between the DCIS and the immune infiltrate areas. This observation suggests that the most pronounced changes in cell type composition occurs at the boundary of these two regions, potentially offering insights into the tumor microenvironment and the interface of cancerous and immune responses. As evidenced in Extended Data Fig. 3, StarTrail also works well on other cell types.

**Figure 3.**
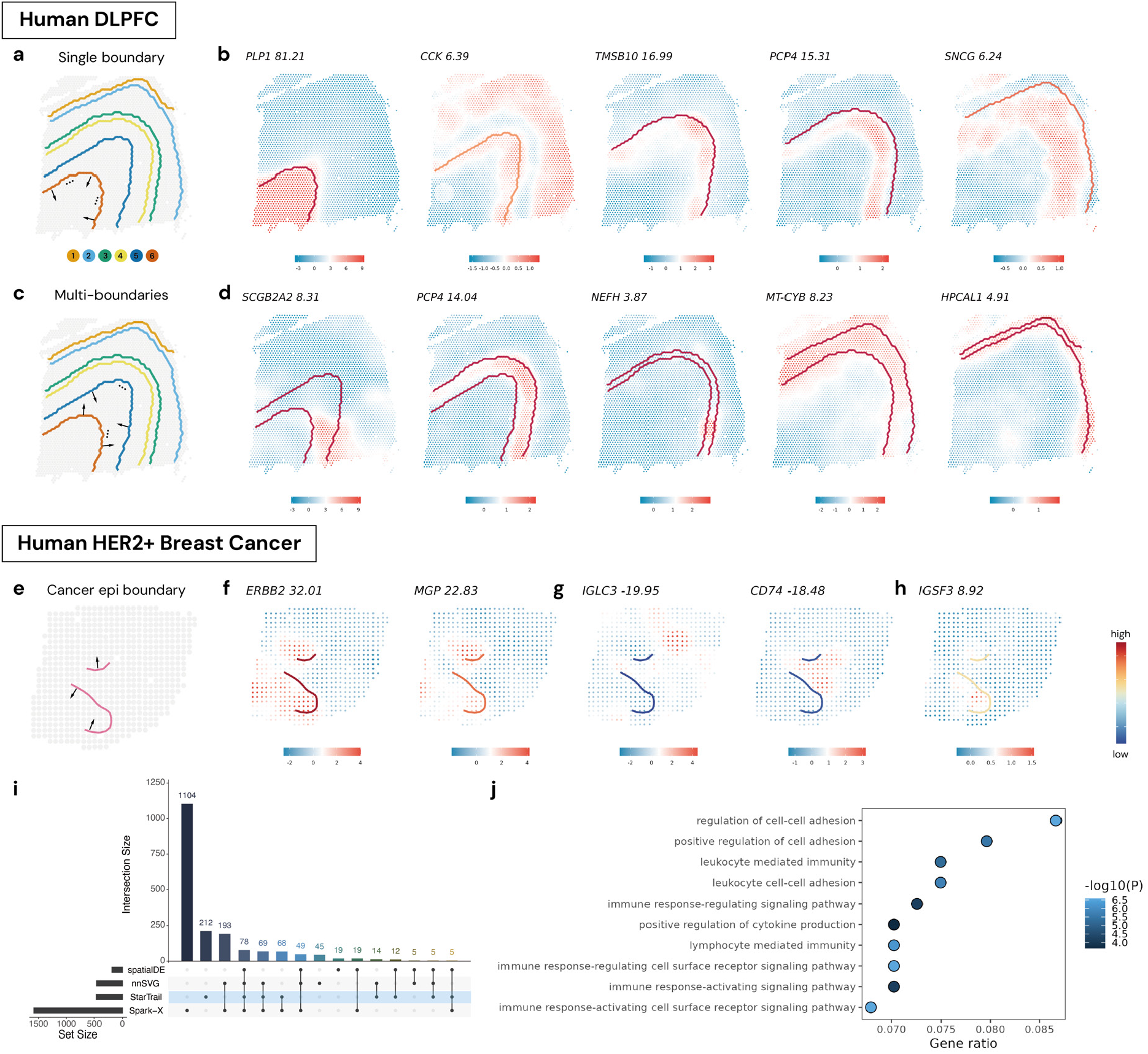
Wombling analysis. **DLPFC**: (a) Annotation boundaries. The arrows indicate the normal direction for a single boundary: between white matter (WM) and layer 6 (L6). (b) Selected genes with top-tier gradients over single boundaries. (c) An example for the normal direction when combining two boundaries: one between WM and L6 and the other between layer 5 (L5) and L6. (d) Selected genes with top-tier weighted-sum gradients over multi-boundaries. **HER2+ breast cancer**: (e) Cancer epithelial cell boundaries. The arrows indicate the normal direction across the boundaries. (f) Top 2 genes with the highest gradients over the two cancer epithelial cell boundaries combined. (g) Top 2 genes with the lowest gradients over the multi-boundaries. (h) Top gene with the highest gradient over the multi-boundaries only detected by StarTrail. (i) Upset plot for detected cliff genes and SVGs. (j) Top 10 pathways identified by cliff genes with the largest gene ratio. All the gene expression in the figure are smoothed and normalized. epi: epithelial.

StarTrail was further tested on spatial proteomic data from human tonsils measured by the CODEX platform [36] (Fig. 2g, Extended Data Fig. 4). The spatial pattern of the tonsil tissue is characterized by the presence of germinal centers (GC), known for heightened abundance of CD15 and a predominance of B cells. StarTrail successfully delineated spatial gradients within this complex tissue structure. The highest *L*^2^ values are indicative of the steepest expression changes, occurring at the GC boundary. Notably, this occurs at the transition zones between two types of B cells.

**Figure 4.**
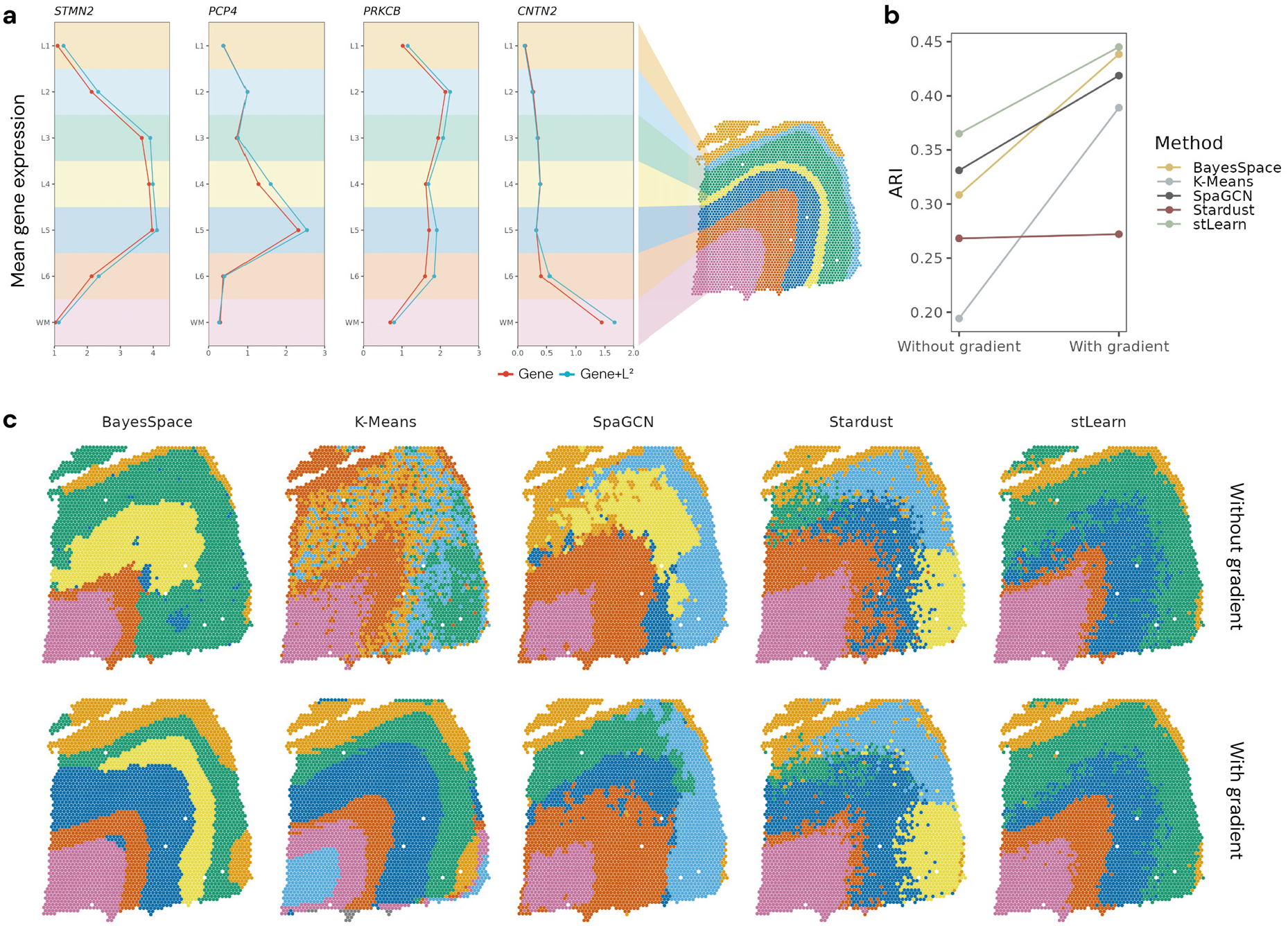
Enhanced clustering analysis. (a) Integrating *L*^2^ to gene expression enlarges the difference of mean gene expression between its enriched layer and other layers. (b) Integrating *L*^2^ to clustering analysis enhanced clustering, shown by higher ARI. (c) The clustering result from five methods. Here, ‘Without gradient’ (the first row) represents the best clustering result (highest ARI) using only gene expression in SVG and cliff gene sets. ‘With gradient’ (the second row) represents the best clustering result using either *L*^2^ alone, or Gene+*L*^2^ in SVG and cliff gene sets. We matched the color of predicted clusters to truth for visualization. See Supplementary Fig. 11 for raw result.

Our results demonstrate that, across various datasets, StarTrail can precisely estimate spatial gradients and delineate areas of rapid changes. StarTrail’s ability to handle multiple data types makes it a unifying tool for integrated omics analysis. StarTrail thus boasts great potential to facilitate the integrative analysis of multiple assays, such as dual gene expression studies or joint analyses of gene expression and proteomics with either computationally inferred or pathological annotations. For example, we could calculate the gradients of one gene and compare that to the gradients of the cell type proportion on the same slide. This adaptability is essential to the current landscape of biological research, where the integration of different omics is increasingly crucial for a deeper understanding of complex biological patterns and dynamics.

### 2.3 Spatial Wombling enables the identification of cliff genes

One of the primary aims in spatial omics research is to illuminate variations between different tissue regions. Based on histology images, pathologists often mark biologically relevant boundaries, which is a labor-intensive process. In the absence of such annotations, computational methods, such as image segmentation or cell type borders identification, can infer such information. These boundaries between tissue regions represent not merely physical divisions, but also transition zones where critical biological changes occur, such as the shift from healthy to diseased tissue. Investigating these boundaries is key to understanding the underlying causes and mechanisms behind the formation of distinct regions, especially in the context of health and disease research, where understanding the factors that influence disease development and progression is critical.

StarTrail leverages these boundaries to identify “cliff genes”, which exhibit sharp expression shifts across boundaries. This cliff-gene approach transcends traditional DE analysis between the two domains separated by the boundaries, by revealing localized gene expression changes that might be missed in DE analysis, where signals are diluted over larger areas. In addition, StarTrail’s capability extends beyond a single boundary. It can adeptly handle multiple boundaries for which traditional DE analyses often struggle due to their intrinsic limitations to two-region comparisons. StarTrail’s flexibility in accommodating different boundary shapes, be they open or closed curves, circumvents the need to assign spots or cells to specific regions, a requirement in DE analysis [16]. This allows for a more dynamic, flexible, and finer-resolution DE and SVG analysis. Focusing on boundary-induced expression changes is essential for understanding the spatial structure of tissues and changes across regions.

#### DLPFC

Delving further into the DLPFC analysis [16], we continued to explore StarTrail’s capabilities to decipher gene expression gradients perpendicular to pathologically annotated boundaries (Fig. 3a). By aggregating these gradients along a single boundary, we were able to identify genes that demonstrate substantial shifts in expression across different regions (Fig. 3b, Extended Data Fig. 5,6, Supplementary Table 1). For example, StarTrail successfully identified *PLP1*, previously noted for pronounced gradients at the WM boundary (boundary 6), as a cliff gene with the highest gradient (*L*^2^ = 81.21) along boundary 6 across all genes. Other genes identified by StarTrail with top-rank gradients along a single boundary also conform to anticipated spatial expression profiles. To validate the accuracy of the estimated gradients, we extended both outward and inward from the pathologist annotated boundaries [37], with each step encompassing one delineated layer of spots (Supplementary Fig. 1,2). For every delineated layer of spots, we computed the average gene expression counts. The concordance between the estimated boundary gradients and the slopes of gene expression change at the boundaries was striking (Pearson correlation between estimated gradient and slope in cliff genes: boundary0: 0.97, boundary1: 0.94, boundary2: 0.99, boundary3: 0.94, boundary4: 0.80, boundary5: 0.95, Extended Data Fig. 7, Supplementary Fig. 2). Genes with a steeper slope across the boundary consistently exhibited higher estimated boundary gradients. This agreement attests to the utility and robustness of our gradient estimation method, confirming that the boundary gradients inferred by StarTrail accurately reflect the actual changes in gene expression as one moves across the pathologically defined boundaries. Gradient estimation along boundaries goes beyond naive exploratory summaries of the change in spatial gene expression. Our boundary-informed supervised analysis thereby facilitates the discovery of “cliff genes” which are intricately linked to biological boundaries.

The biological intricacy of the DLPFC necessitates an analysis that is not confined to single boundary definitions. For example, while *MBP* expression shows a consistent increase from the top right to the bottom left, as indicated by multiple positive boundary gradients, contrasting genes like *PCP4* display heterogeneity in their expression patterns. This is marked by their positive and negative gradients across different single boundaries (Fig. 3b). This variability accentuates the need to accommodate multiple boundaries into spatial omics analyses. When engaging with multiple boundaries, StarTrail aggregates information by taking the weighted sum of gradients along multiple boundaries. For instance, StarTrail enables the delineation of WM-enriched genes through analyzing gradients over boundary 6 (Fig. 3a). Similarly, StarTrail helps to identify genes with high expression in L6 by analyzing the weighted sum of gradients along boundaries 5 and 6 (Fig. 3c). By analyzing multiple boundary pairs (for example, boundaries 5,6 for layer L6, boundaries 4,5 for layer L5, boundaries 3,4 for layer L4, boundaries 2,3 for layer L3, boundaries 1,2 for layer L2), StarTrail accurately identifies genes enriched in specific layers or regions (Fig. 3d, Extended Data Fig. 8).

#### Breast cancer

In addition, StarTrail’s ability to harness estimated gradients for detecting boundaries in various contexts, such as gene expression or cell proportions, extends its utility to the simultaneous modeling of multi-omics data. This capacity is particularly advantageous with co-assay data, for example gene expression and cell type annotation, where cliff genes can be identified in relation to cell-type-defined boundaries. In our study of HER2+ breast cancer, we applied StarTrail, focusing squarely on the previously identified cancer epithelial cell boundary, to estimate boundary gradients for gene expression (Fig. 3e-h, Extended Data Fig. 9, Supplementary Table 2). *ERBB2*, known as a marker gene of cancer epithelial cells, was pinpointed by StarTrail with the highest boundary gradient, demonstrating its efficacy in identifying genes critical to cancer pathology. Further examination of genes with pronounced boundary gradients revealed their biological significance in cancer progression. For instance, *MGP*, whose gene expression was found to be negatively associated with patients’ overall survival, is a promising prognostic biomarker [38]. Similarly, increased *CD24* expression in breast cancer tissues compared to normal tissues, as observed in TCGA, underscores its potential as a potential biomarker for prognosis [39]. When contrasting the top genes identified by cell type-specific boundaries with those discerned by pathologist-annotated cancer boundaries, we observed remarkable concordance (Extended Data Fig. 9). This suggests that, in the absence of pathological annotations, cell type boundaries can serve as an effective proxy for delineating the dynamic interface between cancerous and non-cancerous regions. These results demonstrate StarTrail’s precision in delineating spatial expression boundaries and corroborate the robustness of StarTrail’s gradient estimation techniques. StarTrail’s ability to accurately identify genes with drastic expression changes at cellular boundaries enables researchers to focus on regions of rapid changes, which often harbor critical biological insights. In the context of disease pathology, such as cancer, discovery of these rapidly changing zones can provide a wealth of information regarding the mechanisms of disease progression, invasion, and interaction with the surrounding tissue microenvironment. By targeting these rapidly transitional areas, researchers can gain a more comprehensive understanding of omics dynamics and cellular heterogeneity at the tumor margin, which is essential for the development of targeted therapies and better predictive prognostic markers.

### 2.4 StarTrail-identified cliff genes and pathways offer novel insights

We benchmark StarTrail’s performance in identifying cliff genes against established SVG detection methods in the human DLPFC and HER2+ breast cancer dataset. We chose four widely-used SVG methods: Spark-X [40], SoMDE [41], nnSVG [42], and SpatialDE [43].

#### DLPFC

Analyzing the DLPFC 151576 dataset, among 11,686 genes, Spark-X, SpatialDE, nnSVG, and SoMDE identified 9,673, 5,266, 1,622 and 1,185 SVGs respectively; and StarTrail identified 758 cliff genes (Supplementary Fig. 3, Supplementary Table 1). Notably, in this dataset, most cliff genes detected by StarTrail were identified by some SVG method(s), with only 7 StarTrail-exclusive genes missed by all four SVG methods (see Methods for threshold recommendations). Investigation into these 7 unique genes revealed a commonality: each exhibited rather localized patterns of hot spots along at least one pre-defined boundary (Supplementary Fig. 4). To substantiate the validity of StarTrail findings beyond visual assessments, we checked cliff genes that StarTrail identified from one single slide against genes identified by combining information across 8 slides from the spatialLIBD study [16]. Reassuringly, we observed a substantial overlap: for instance *CCDC80, MEX3D*, and *RRP8* showed significance at the L6-WM boundary, while *MSX1* and *ID4* were significant at the L1-L2 boundary (as detailed in Supplementary Table S4C of the spatialLIBD paper).

#### Breast cancer

we similarly compared cliff genes identified by StarTrail with genes detected by the SVG methods. In this dataset, spatialDE, nnSVG, and Spark-X identified 192, 458, and 1,585 SVGs respectively. StarTrail identified 463 cliff genes at either of the two cancer epithelial cell boundaries, and the weighted-sum of the two boundaries gradients led to 100 cliff genes (Fig. 3i, Supplementary Table 2, Supplementary Fig. 5, 6). Note that SoMDE did not detect any SVGs and was thus omitted from further analysis. Among the 463 (100) cliff genes, 212 (12) were StarTrail-exclusive, not recognized by any SVG method. The 12 genes (Supplementary Fig. 6) manifest high expression in either cancer in situ region (e.g. *IGSF3*) or the immune infiltrate region (e.g. *MBP*). *IGSF3* (Fig. 3h), exhibiting the largest gradient in StarTrail’s multi-boundaries analysis but undetected by all SVG methods, is part of the immunoglobulin superfamily (IgSF), which includes members whose gene expression levels are known to vary in breast cancer, making them potential prognostic biomarkers. [44]. *IGSF3*, in particular, has been highlighted as a possible target for Chimeric antigen receptor (CAR) therapy and is considered essential in 80% of head and neck squamous cell carcinoma cell lines (classified as “common-essential” by DepMap) [45]. Moreover, TCGA DE analysis between tumor and adjacent normal tissue revealed significant differences in over 90% of IgSF genes in at least one cancer type. Specifically, *IGSF3* demonstrates a marked over-expression in breast invasive carcinoma samples from TCGA, highlighting its relevance and potential impact in the context of breast cancer [46].

StarTrail not only identifies cliff genes that offer insights missed by existing methods but also uncovers undetected pathways. Gene Ontology (GO) analysis was conducted to delve into the pathways of these cliff genes (Fig. 3g) [47]. Cliff genes identified at either boundary revealed 54 pathways missed by SVG methods; and using cliff genes detected with the multi-boundaries approach revealed 30 missed pathways (Supplementary Tables 3-8). Notably, GO results highlight terms pertinent to the response to interleukin-1 (IL-1, GO:0071347 and GO:0070555, Supplementary Fig. 7), one of the major pro-inflammatory cytokines known to be elevated in various tumor types including breast cancer. IL-1 has been linked to tumor progression through its role in promoting the expression of genes involved in metastasis, angiogenesis, and growth factors [48]. Among the 30 GO terms uncovered from StarTrail’s multi-boundaries approach, multiple are related to immune response, including the regulation of natural killer cell mediated immunity (GO:0002715, Supplementary Fig. 8) and the positive regulation of myeloid leukocyte mediated immunity (GO:0002888), presenting granular insights into the immune dynamics within the tumor microenvironment. As StarTrail is designed to detect localized spatial patterns, it holds the promise of offering superior power and resolution in scenarios where multiple unconnected hot spots occur around boundaries, uncovering localized patterns that may well be overlooked by traditional SVG detection methods.

So far we have concentrated on the absolute magnitudes of the gradients. However, the gradients’ sign can provide orthogonal insights. Focusing on cliff genes characterized by a negative gradient (indicating lower expression within the cancerous regions), we observed a strong association with immune-related processes (Supplementary Table 5). Among the top 15 significant pathways identified by StarTrail, an impressive 14 were directly tied to immunity, antigens, and B cell functions. This starkly contrasts with the findings from SVG detection methods, where nnSVG, Spark-X, and SpatialDE identified 4, 3 and 6 pathways related to immunity.

### 2.5 Enhanced downstream analysis with StarTrail inferred finer spatial dynamics

In addition to StarTrail’s direct benefits of precisely demarcating boundaries and detecting cliff genes, integrating StarTrail inferred gradient information can significantly enhance the inferential capabilities of traditional spatial methodology. Although a complete evaluation is beyond the scope of this work, we demonstrate in this section how harnessing the output of StarTrail inferred spatial dynamics can bolster spatial domain detection, commonly referred to as clustering analysis.

It is important to note that we are not introducing a new clustering method. Rather, we performed integration by feeding StarTrail inferred spatial dynamics as input to existing clustering methods. We propose two approaches for integration: first, using the *L*^2^ norm of gradients as the input for clustering, and second, combining the original omics data with the *L*^2^ norm of gradients as input. Since high *L*^2^ values typically correspond to areas of rapid omics profile change, which often corresponds to biological boundaries, the inclusion of *L*^2^ enhances the distinction between different spatial regions. To illustrate this, we show four genes with diverse expression patterns (Fig. 4a). When *L*^2^ values are included with gene expression, the contrast between areas enriched for each gene becomes more pronounced.

We applied several clustering algorithms, including both traditional clustering methods like K-means [49, 50] and spatial clustering methods: BayesSpace [12], SpaGCN [11], Stardust [14] and stLearn [13], to the DLPFC dataset based on gene expression alone, *L*^2^ norm alone, and the sum of gene expression and *L*^2^ (Gene+*L*^2^). In our analysis, we used two sets of genes: highly variable genes (selected by SPARK [9]) and cliff gene sets (genes with absolute boundary gradients greater than 5 at any boundary). The choice of gene set (either SVGs or cliff genes) and input metric(s) can both influence clustering outcomes, with no universally optimal selection for all methods. However, when we compared the best Adjusted Rand Index (ARI) [51] scores for clustering methods with and without gradients, the inclusion of StarTrail inferred gradients consistently outperformed analyses relying solely on gene expression (Fig. 4b,c, Extended Data Fig. 10, Supplementary Fig. 11). Notably, K-Means clustering with *L*^2^ demonstrated a marked improvement, revealing a distinct and drastically clearer layer structure (Fig. 4b,c).

These findings underscore the potential of including StarTrail inferred spatial dynamics to enhance the resolution and accuracy of clustering analyses in spatial omics studies, offering a more comprehensive and granular understanding of tissue architecture and function.

## 3 Discussion

StarTrail represents a pioneering and paradigm-shifting approach in the spatial omics field as the first method that employs a rigorous yet efficient spatial gradient framework to empower inference at highly localized, potentially disjoint regions that existing methods have little power. This innovative technique substantially enhances our understanding of spatial omics feature by providing directional, quantitative, and high-resolution gradient (i.e., rate of change) information. Augmenting the original omics measurements with such gradient information, StarTrail sheds new light on tissue structure and function.

The fundamental strength of StarTrail lies in its utilization of gradient information, which has multiple advantages in practical applications. The gradient flow map, a central feature of StarTrail, provides an intuitive visualization of omics dynamics, revealing the direction of changes across tissue samples. This is particularly valuable in identifying areas of rapid transition within the tissue, thereby aiding in the detection of critical biological boundaries. Another significant application is the identification of “cliff genes” – genes that exhibit dramatic shifts in expression at specific tissue interfaces. This ability to pinpoint genes that are highly variable across spatial boundaries has profound implications beyond the routine analysis of SVGs. Furthermore, StarTrail is specifically tailored to precisely demarcate local boundaries and to detect local omics patterns near pre-defined boundaries. It is therefore complementary to SVG methods proposed in the literature. For instance, significant SVGs with small gradient values at any boundary suggest alternative boundaries unknown to us, or the presence of potential new clusters (Supplementary Note, Supplementary Figs. 9, 10). For downstream analysis such as clustering analysis, the inclusion of gradient information significantly enhances the resolution and accuracy of identifying distinct spatial domains within tissue samples.

Another pivotal advantage of StarTrail is its computational efficiency. Utilizing NNGP models, Star-Trail dramatically reduces the computational cost required for analysis – from 40 hours to a mere 2 minutes per gene per slide, as evidenced in our analysis of the DLPFC dataset. This efficiency is achieved without sacrificing accuracy, making StarTrail a practical tool for large-scale studies. Furthermore, Star-Trail’s flexibility in handling multiple data types – from measured transcriptomics or proteomics data to computationally inferred annotations – makes it a versatile tool for various spatial omics applications and the analysis of co-assays.

Looking forward, there are several promising directions for extending StarTrail’s capabilities. For example, the development of spatial gradient processes that model multiple omics features simultaneously is highly warranted. This could provide a more comprehensive understanding of the complex interactions among multiple genes or proteins within the same spatial context. Another promising direction is the development of clustering methods that are specially designed to incorporate gradient information. Such methods could offer a more powerful approach to segmenting tissues based on dynamic changes, leading to more accurate and biologically relevant clustering outcomes.

In conclusion, the introduction of StarTrail to the spatial omics toolkit has the potential to open doors to a deeper understanding of the spatial dynamics of biological tissues. This novel approach promises to enhance our comprehension of tissue structure and function by leveraging the rich information contained within spatial gradients.

## Acknowledgments

DL was supported by NIH grants R01 AG079291, R56 LM013784, R01 HL149683 and UM1 TR004406. JC, GPG, YL are supported by a Department of Defense Breast Cancer Research Program Award (HT94252310961). JC, QS and YL are supported by R01 AG079291, U01 HG011720, U01 DA052713, and P50 HD103573.

## 4 Method

### The StarTrail pipeline

The analysis in StarTrail begins by inputting data from spatial omics experiments or its annotation. For omics data, such as gene expression or protein abundance, we first implement a smoothing step to enhance signal quality and reduce noise. The omics measurement of each spot/cell is smoothed as the weighted sum of its neighbors and itself. This is followed by data normalization using the SCTransform function in Seurat [52]. For cell type proporiton, only the smoothing step is applied. This initial phase refines raw data, preparing them for subsequent analysis.

The core of StarTrail’s workflow involves employing the NNGP [29] [30]. Here, we model the relationship between coordinates *s* and omics feature (e.g., gene expression) *Y* using the following Bayesian hierarchical model [53] [54]

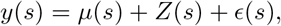

where *μ* is a mean function, *Z* ∼ GP(0, *K*) is a zero mean GP with covariance function *K*, and *ϵ*(*s*) ∼ *N* (0, *τ* ^2^) is a zero mean white-noise process capturing measurement error, also known as nuggets. In particular, the GP *Z* is a stochastic process with finite d imensional r ealization b eing multivariate Gaussian: for any *s*_1_, …, *s*_*n*_ ∈ ℝ^2^,

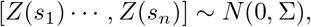

where ∑_*ij*_ = *K*(*s*_*i*_, *s*_*j*_) is the covariance matrix calculated based on predefined kernels. Here we choose the Matérn kernel as the covariance function due to its flexibility in controlling smoothness [55]:

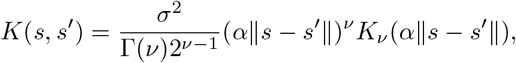

where *K*_*ν*_ is the modified Bessel function of the second kind.

Traditional GP modeling is computationally intensive, primarily due to the inverse of the *n* by *n* covariance matrix ∑, at a cost of *O*(*n*^3^). StarTrail, adopting the NNGP approach, significantly improves efficiency and scalability, crucial for handling large datasets common in spatial omics studies. NNGP, by focusing on the *m*-nearest neighbors and a sparse approximation to the covariance, reduces the computational cost to *O*(*nm*^3^) ≪ *O*(*n*^3^) [29] [30]. Here we chose *m* = 10.

We implemented the spNNGP function (“response” algorithm) in the spNNGP package [30]. The priors for the kernel parameters and additional tuning parameters are available in our GITHUB repository.

After fitting the model, the next step is gradient estimation. In the traditional spatial gradient process, the gradient ∇*Z* and curvature ∇^2^*Z* are estimated by inferring joint distribution of [*Z*, ∇*Z*, ∇^2^*Z*] with cross-covariance characterized by

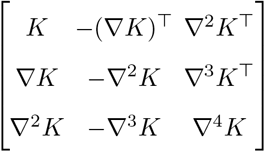

Given the Matérn kernel is ⌈*ν*⌉ − 1 times mean square differentiable (MSD) [55], we set the prior of *ν* to be supported on (2, 5) to ensure that the GP with kernel *K* is 2-times MSD, i.e., ∇^2^*Z* is well-defined. Due to the nature of NNGP, and because we do not fix *ν* in the kernel, the closed form of ∇*K* is not available. As a result, we propose to estimate the gradients using finite differences [56], a mathematical method for approximating derivatives. Here, we explain the case for 2D coordinates, calculating the gradient in the direction of *e*_1_ = (1, 0) and *e*_2_ = (0, 1). We denote the posterior mean of *f* as 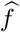, and for any given location *s*, 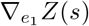 and 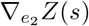 are approximated by

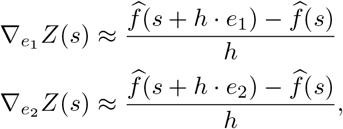

where *h* is a sufficiently small step size specified by users. According to our empirical observation, we recommended and set as default *h* = 0.8*ι*, where *ι* = min_*i,j*_ ∥*s*_*i*_ − *s*_*j*_∥ is known as the minimal separation of the locations 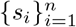 .

Similarly, the curvatures at *s* are approximated by:

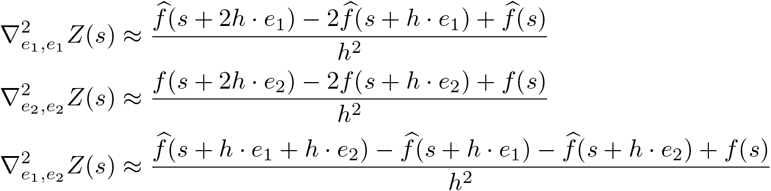

Consequently, the gradient at *s* can be estimated by 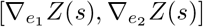.

Wombling analysis in StarTrail calculates gradients along a given curve *γ* : [0, *T*] → ℝ^2^, taking segment points 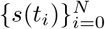 as input and adopting a piece-wise linear approximation. For each line segment *γ*(*t*) with starting point 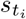 and ending point 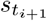, let 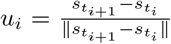 be the unit vector representing this segment. Then the normal unit vector *v*_*i*_ = *Ru*_*i*_ with 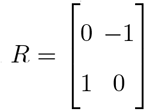 being the rotation matrix of 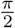.

An approximate to 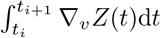 is given by the Simpson’s rule [57][58]

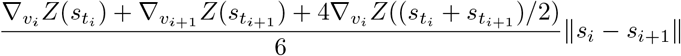

where

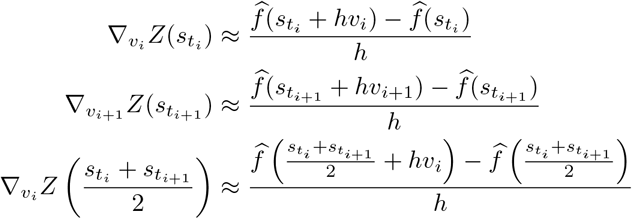

Then the average gradient along the curve *γ* is approximated as

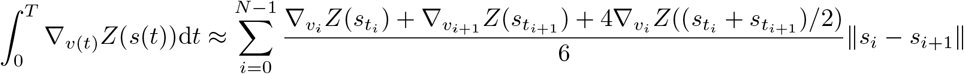

For easier comparison across curves, we scaled the approximated gradient of each curve using its piece-wise linearly approximated length 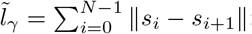:

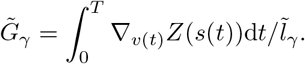

When considering multiple boundaries *γ*_1_, …, *γ*_*k*_, the weighted-sum gradient is approximated as

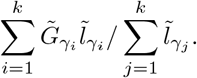

### Boundary detection in StarTrail

StarTrail excels in precisely detecting boundaries within spatial omics data. This process begins by identifying spots or cells that exhibit high *L*^2^, specifically those exceeding a predefined quantile threshold (default is 0.9). Subsequently, the dbSCAN algorithm [59] is deployed to cluster these points, creating an initial map of areas where substantial changes coalesce, and thus suggesting potential boundary locations. For each cluster identified by dbSCAN, a principal curve is fitted. This curve acts as a smooth line that passes through the center of each cluster, effectively delineating the boundary. The default number of nodes in each principal curve is set as the number of points in the cluster divided by ten. Through this methodical approach, StarTrail offers a nuanced and precise way to detect boundaries within spatial omics data. By combining rigorous statistical techniques with advanced clustering algorithms, StarTrail effectively highlights regions of rapid changes, providing valuable insights into the spatial organization and underlying biological dynamics of the sampled tissue.

### Cliff gene gradient threshold

For DLPFC, we used 0.25*σ*_0.9_*/ι* as the threshold, where. *σ*_0.9_ is the 0.9 quantile of the standard deviation of smoothed gene expression, and ι is the minimal separation between spots. 0.25 here accounts for the change spanning across two rows/columns of spots over half of the given boundary. For HER2+ analysis, we used 0.5*σ*_0.9ι_. We note here that this threshold reflects the actual change of gene expression and is therefore not a probability. Thus, users could adjust this threshold based on their needs.

### Gene+*L*^2^

Gene+*L*^2^ is designed to augment the original gene expression data with a gradual spatial change information. To avoid overshadowing the inherent characteristics of gene expression, we meticulously regulate the magnitude of the *L*^2^ term. Specifically, we use a scaled *L*^2^ (divided by the range max *L*^2^ − min *L*^2^). This approach ensures that the alteration to each value does not exceed 1, preserving the nuanced nature of the original data.

### Simulations

To mimic a spectrum of patterns in ST data, we simulated data under several distinct settings. For simulation 1 (Extended Data Fig. 1 top row), we simulated data from trigonometric functions:

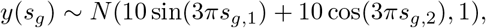

where *s*_*g*,1_ and *s*_*g*,2_ are vectors spaced equally from 0 to 1 in increments of 0.05. In the second simulation (Extended Data Fig. 1 second row), we created a linear gradient pattern, or ‘streak’, where the data transitions gradually from a background level to a higher level towards the center in the (0, 1) direction. The third simulation involved the creation of a ‘hot-spot’ pattern (Extended Data Fig. 1 third row), which similarly sees a gradual change from the background parameter to a center of a circle. The fourth simulation produced a ‘layer’ pattern (Extended Data Fig. 1 bottom row), characterized by a sharp transition from one level to another, resembling discrete layers. All the simulated data can be accessed in our GitHub repository.

### Clustering analysis

We performed clustering with five methods on gene expression, *L*^2^, or Gene+*L*^2^ to cluster spatial locations, using SVGs (top 3000 SVGs selected by SPARK[9]) or cliff gene sets (absolute gradient at any boundary greater than 5). The five methods include BayesSpace [12], K-Means [49, 50], SpaGCN [11], Stardust [14] and stLearn [13]. BayesSpace [12], a fully Bayesian method, leverages spatial neighborhoods information to enhance resolution in ST data and perform clustering. K-Means [60] partitions a dataset into K distinct, non-overlapping clusters by minimizing the variance within each cluster. SpaGCN [11] employs a graph convolutional network to identify SVGs and spatial domains. Stardust [14] integrates gene expression with spatial location to create a pairwise distance matrix for clustering using the Louvain algorithm [61]. stLearn [13] utilizes Spatial Morphological gene Expression normalization (SME normalization) based on gene expression, spatial location, and image data, followed by K-Means clustering [60]. For each method, we adopted the standard pipelines and followed their recommended parameter settings. For BayesSpace, SpaGCN, Stardust, and stLearn, we applied log-transformation and Principal Component Analysis (PCA) to all three types of data (i.e., gene expression, spatial location, and image data). We did not perform any transformations on the input for K-Means. For Gene+*L*^2^ in BayesSpace, we add rounded and scaled *L*^2^ to the gene expression matrix, as the developers recommended when the data contain values between 0 and 1. In BayesSpace, K-Means, SpaGCN, and stLearn, the number of clusters is set based on the ground truth (here we used 7 for DLPFC 151676). Stardust does not have the option to specify the number of clusters.

### De-convolution analysis

We applied RCTD for cell type de-convolution to the HER2+ breast cancer data. RCTD is a computational technique that uses single-cell RNA-seq data to decompose cell type mixtures while adjusting for technical variations. We set the doublet mode to be “full” to obtain the proportion of all the cell types in each spot.

### Gene ontology analysis

We performed gene ontology (GO) analysis using R package GO.db. We included three GO categories, molecular function (MF), biological process (BP) and cellular component (CC).

## Data availability

The DLPFC [16] data is obtained from https://research.libd.org/globus/. The breast cancer data are obtained from https://doi.org/10.5281/zenodo.4751624 (ST) and https://singlecell.broadinstitute.org/single_cell/study/SCP1039 (scRNA-seq reference). The CODEX [36] data is shared by the MaxFuse [37] authors.

## Code availability

All code used in this study, including the StarTrail software, can be found at https://github.com/JiawenChenn/StarTrail.

## 5 Extended Data

**Extended Data Fig. 1.**
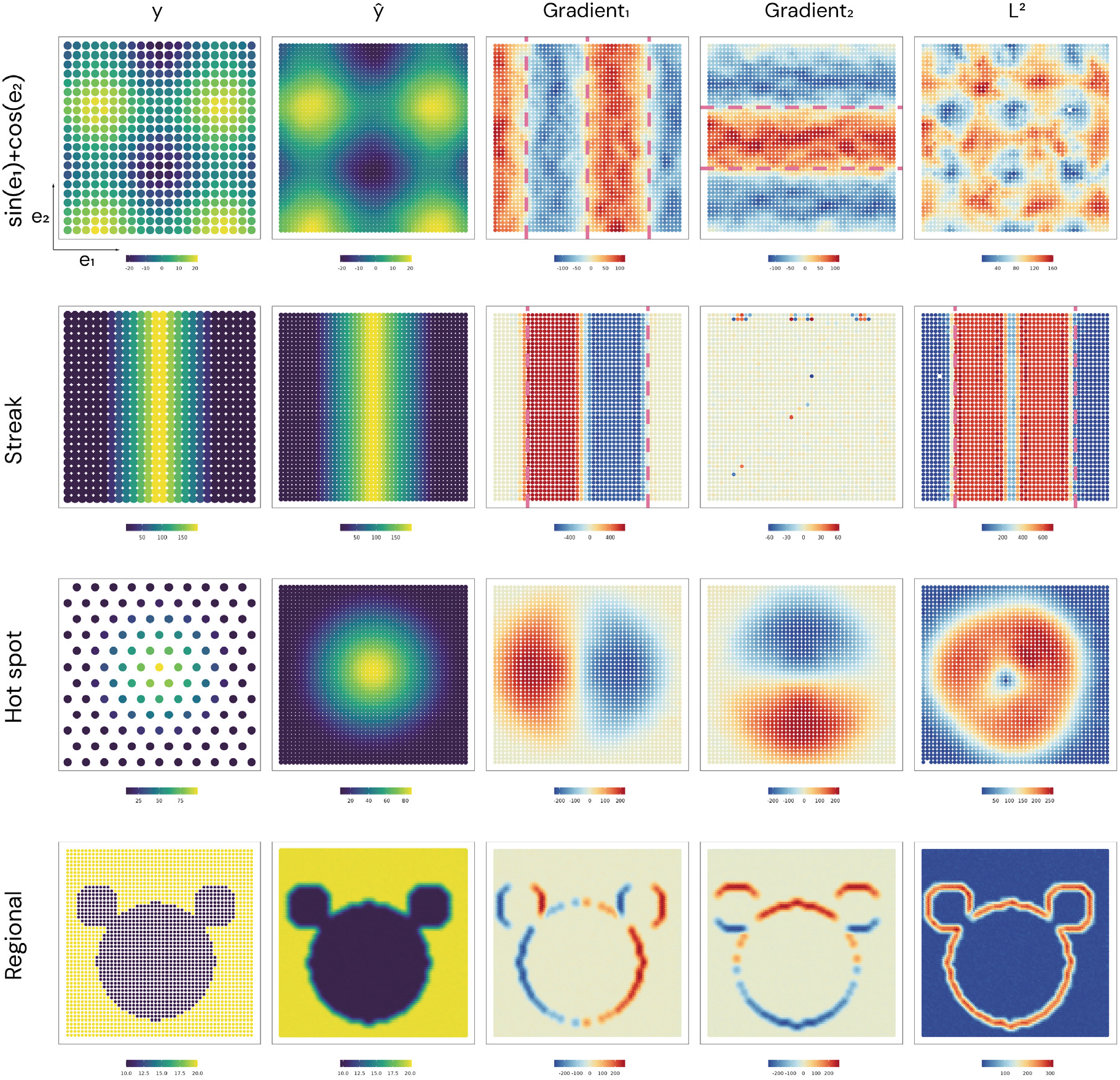
Simulation study. Measurements are simulated from a mixture of sine and cosine functions, streak, hot spot and regional patterns.

**Extended Data Fig. 2.**
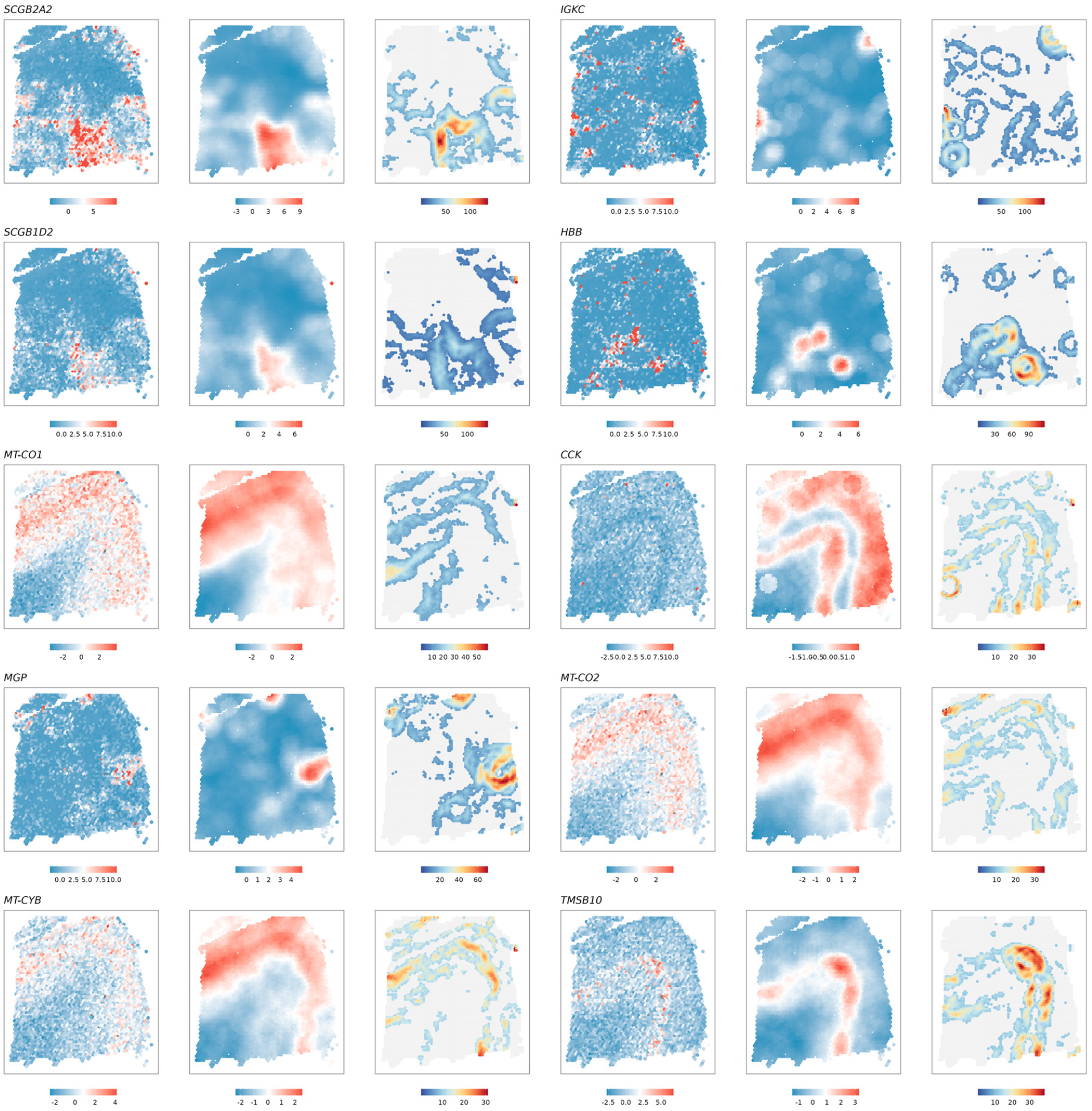
Estimated gradients for gene with top 10 median *L*^2^ values in DLPFC. For each gene, the left panel is the normalized gene expression, the middle panel is the smoothed and normalized gene expression, and the right panel is the estimated *L*^2^. Spots with *L*^2^ smaller than 0.7 quantile are masked by gray color.

**Extended Data Fig. 3.**
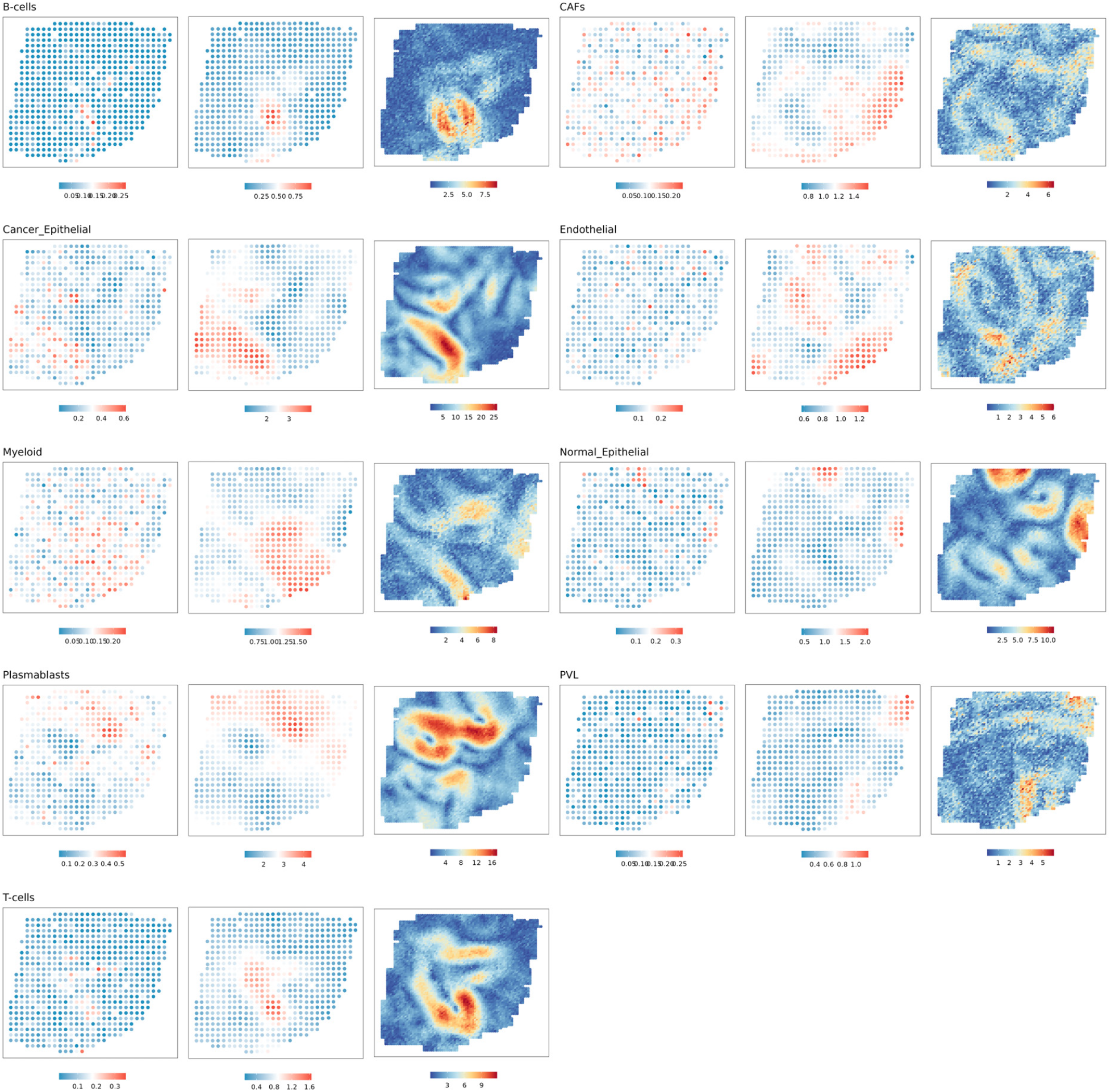
Estimated gradients for cell type proportion in HER2+ breast cancer data. For each gene, the left panel is the cell type proportion, the middle panel is the smoothed cell type proportion, and the right panel is the estimated *L*^2^. Spots with *L*^2^ smaller than 0.7 quantile are masked by gray color.

**Extended Data Fig. 4.**
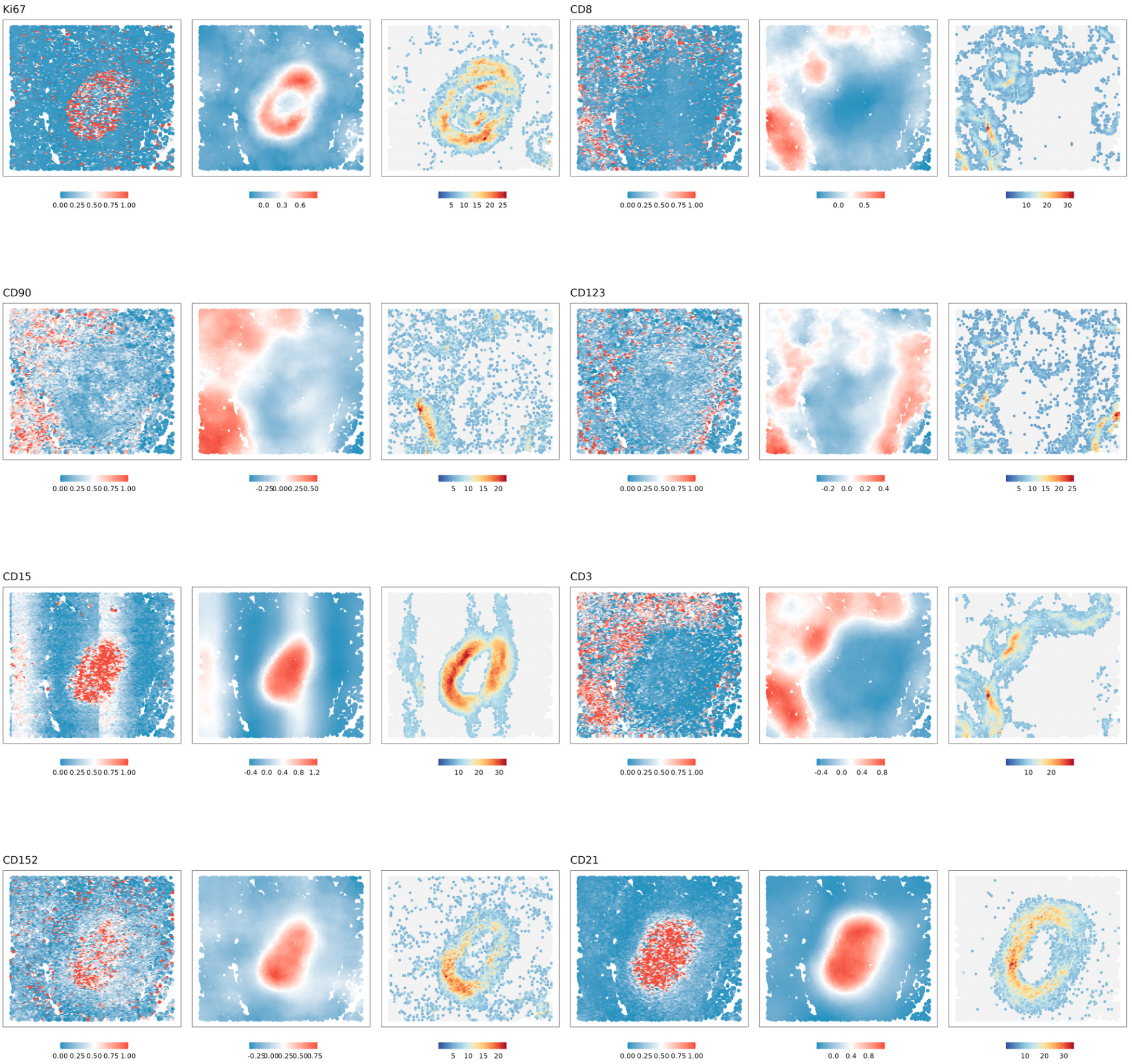
Estimated gradients for selected proteins in CODEX data. For each protein, the left panel is the normalized abundance, the middle panel is the smoothed and normalized abundance, and the right panel is the estimated *L*^2^. Spots with *L*^2^ smaller than 0.7 quantile are masked by gray color.

**Extended Data Fig. 5.**
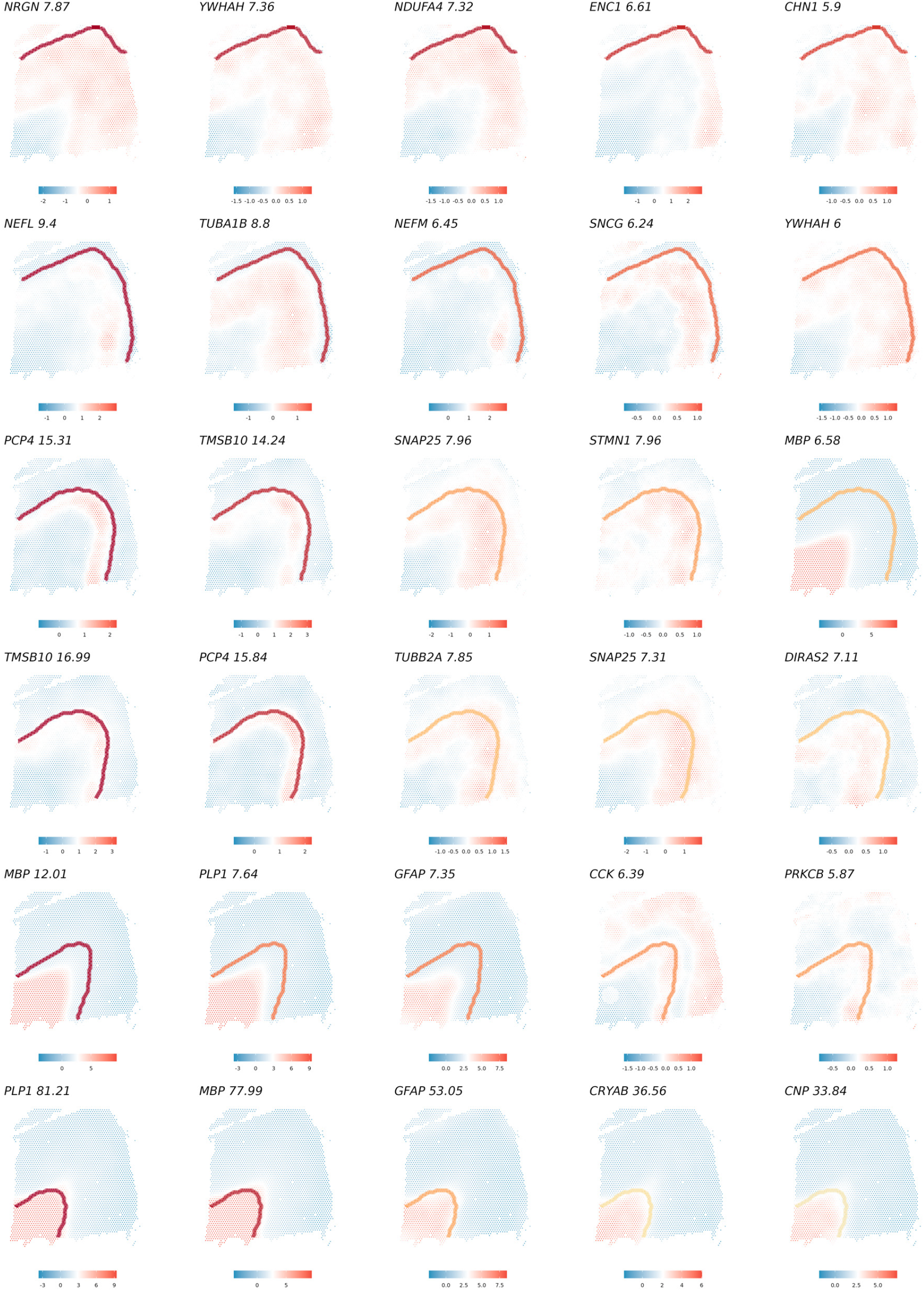
Cliff genes with top five positive gradient for each boundary in DLPFC.

**Extended Data Fig. 6.**
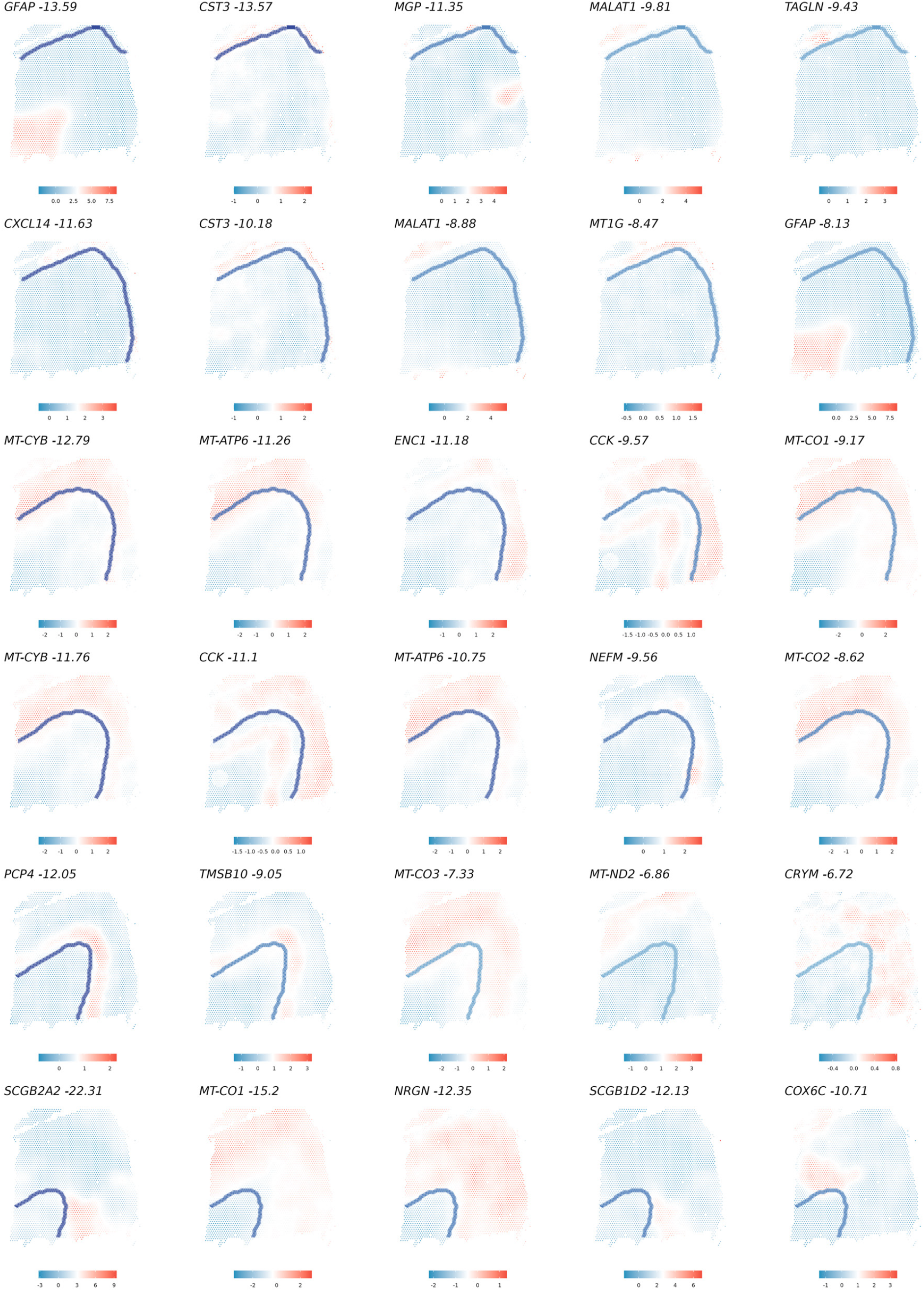
Cliff genes with top five negative gradient for each boundary in DLPFC.

**Extended Data Fig. 7.**
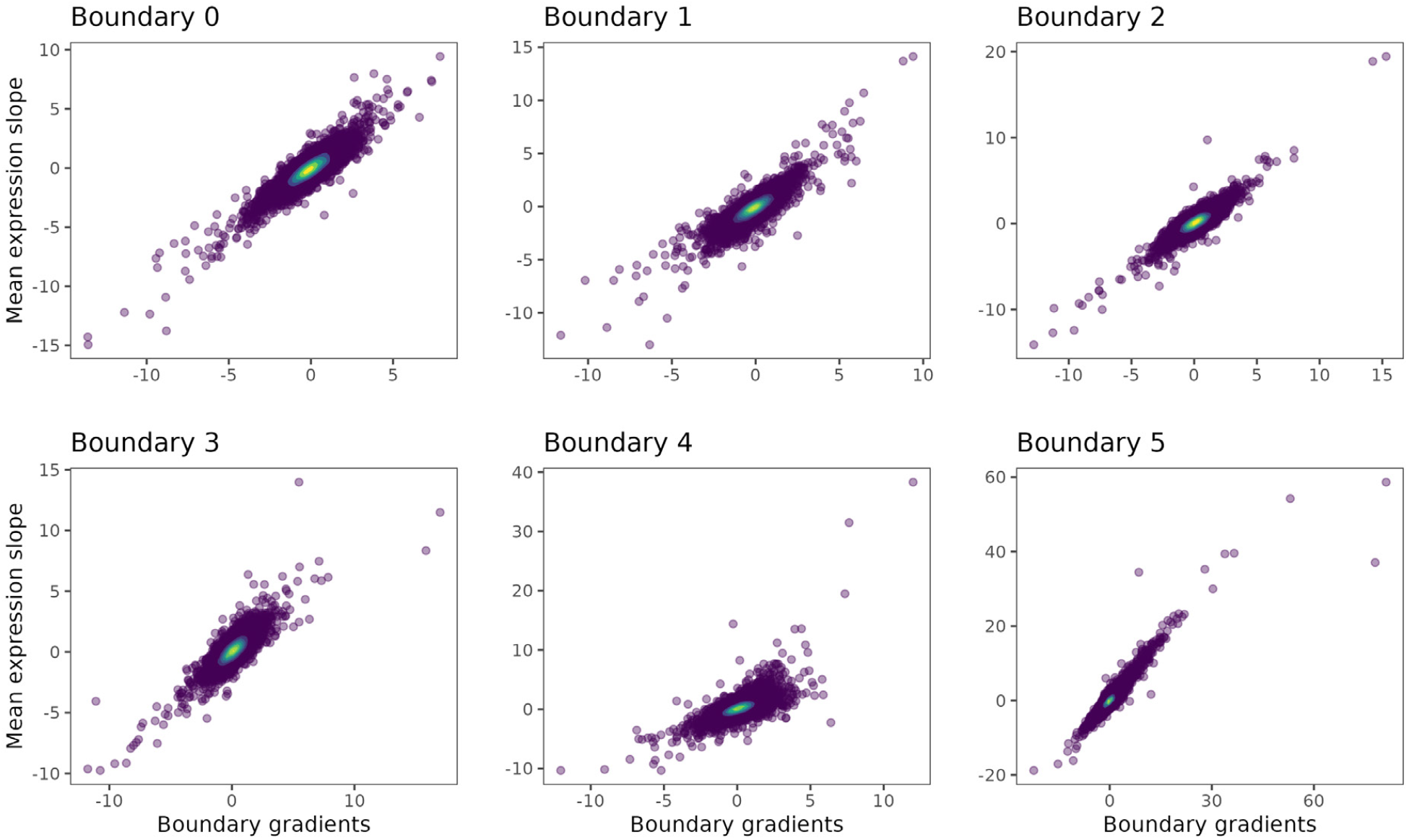
Comparison between mean expression change slope (spots in distance 0 layer to -0.05 layer) and estimated boundary gradients.

**Extended Data Fig. 8.**
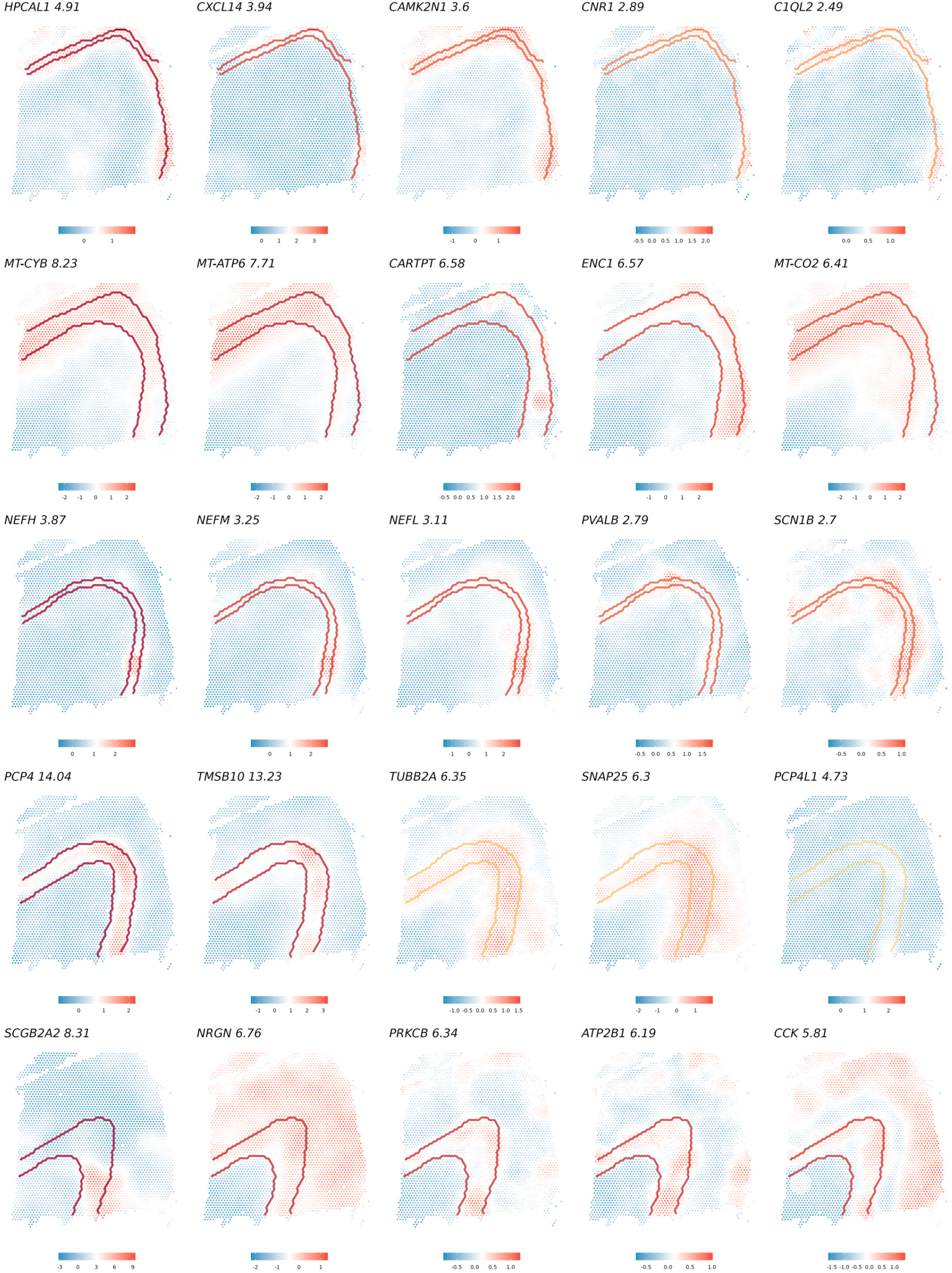
Cliff genes with top five positive gradient for each boundary pair in DLPFC.

**Extended Data Fig. 9.**
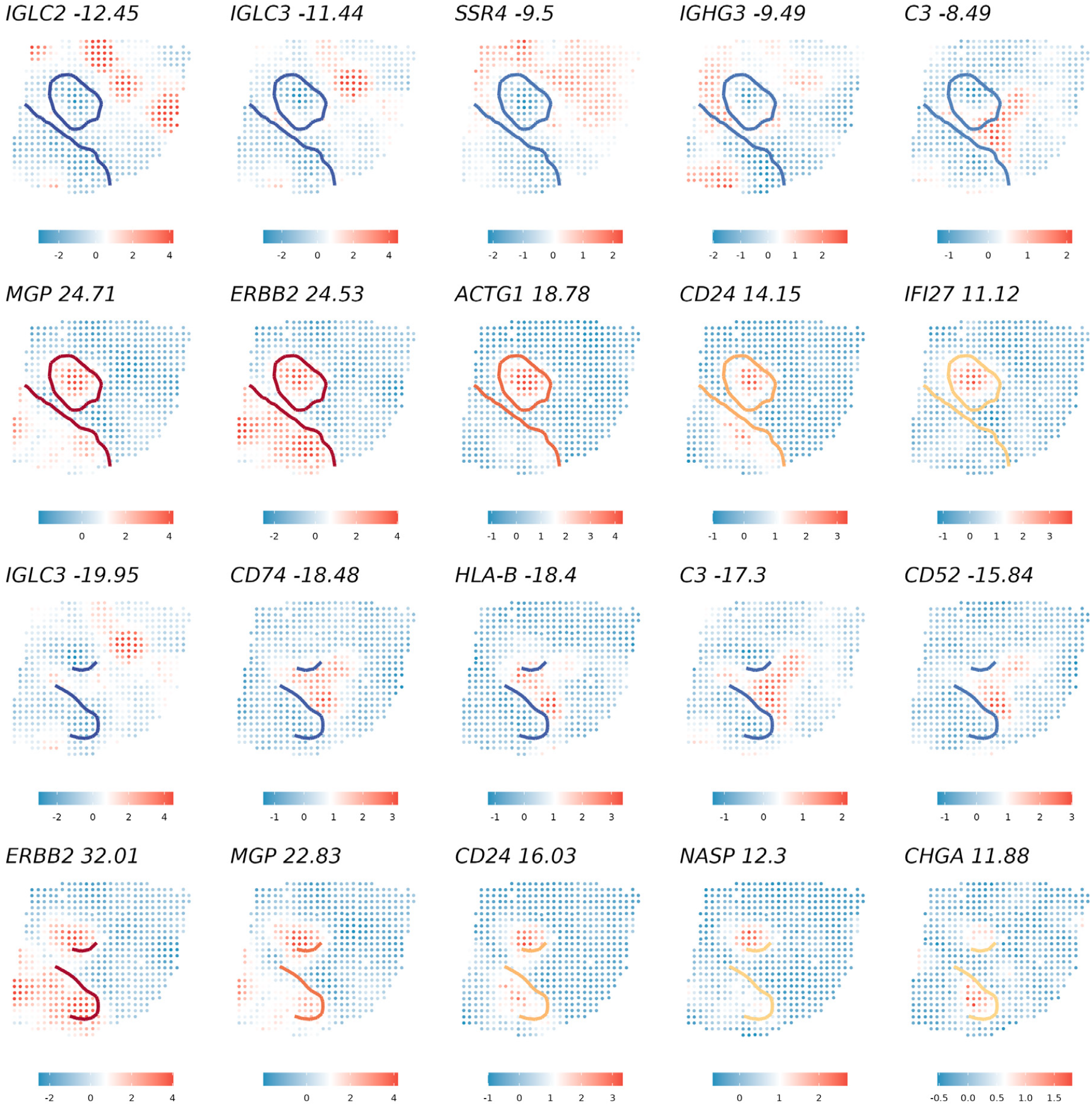
Cliff genes with top five gradient for multi-boundaries in HER2+ breast cancer. (Row 1) top five genes with positive gradients for hand-drawn cancerous boundary based on pathologist annotation. (row 2) top five genes with negative gradients for hand-drawn cancerous boundary based on pathologist annotation. (Row 3) top five genes with negative gradients for cancel epithelial cell boundaries. (Row 4) top five genes with positive gradients for cancel epithelial cell boundaries.

**Extended Data Fig. 10.**
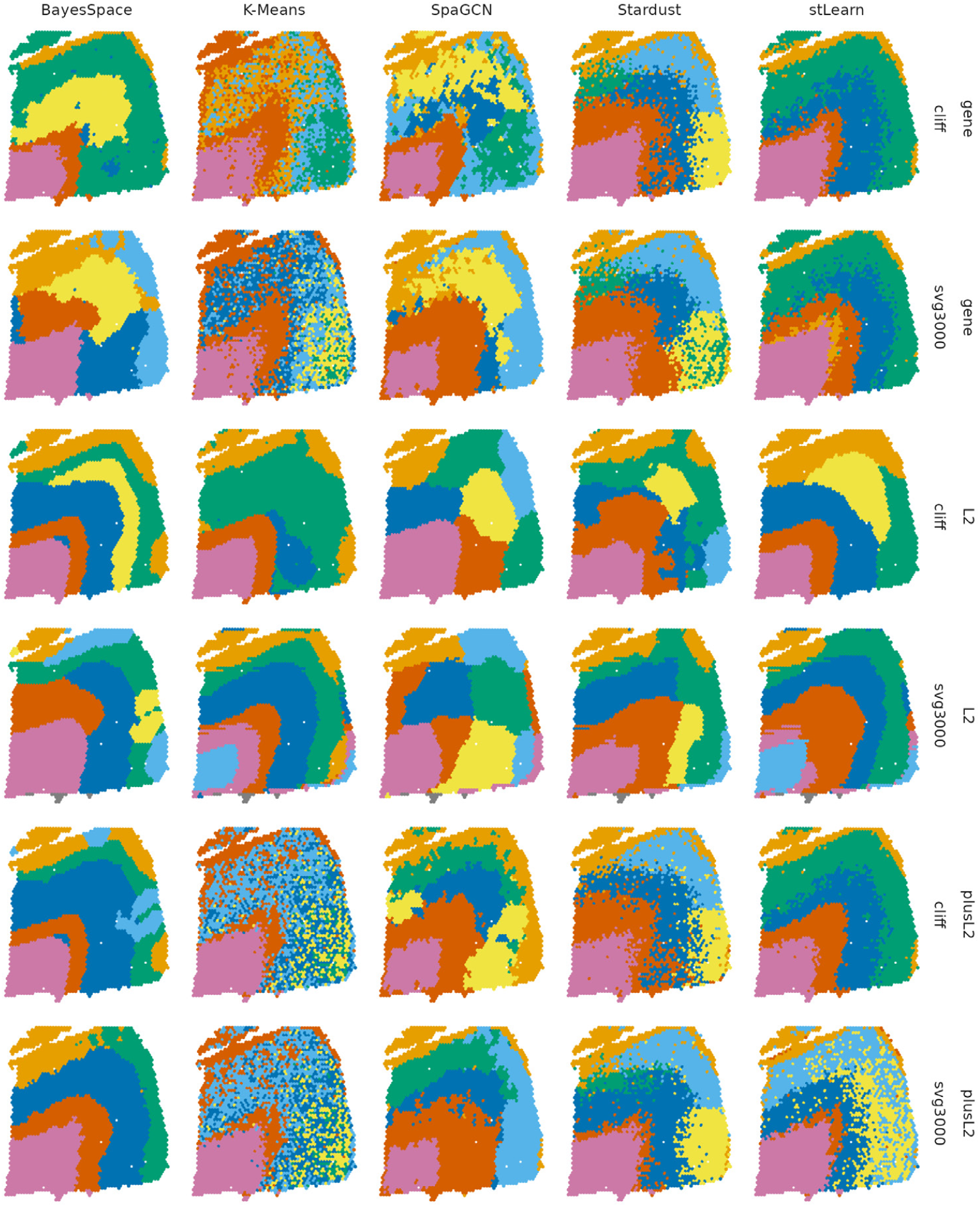
Refined clustering results for all methods with different input (gene expression, *L*^2^, Gene+*L*^2^) in different gene sets (HVG, cliff genes).

## Supplementary Note for “Investigating spatial dynamics in spatial omics data with StarTrail”

### 1 Analysis of non-cliff SVGs

StarTrail, used together with SVG methods, can reveal novel biological insights. For the DLPFC dataset, upon examining significant SVGs with minimal gradient values, it becomes apparent that many genes exhibit expression changes across multiple boundaries (Supplementary Fig. 9). Notably, genes such as *HMGCR* and *BOLA3* manifest high expression in the bottom right of the slide, indicative of potentially new subregions not yet recognized in literature. These subregions potentially account for the distinct cluster observed in the bottom right region in various clustering analysis results (as detailed in Supplementary Fig. 12 of the spatialPCA paper [62]).

In the analysis of HER2+ breast cancer, we discovered that certain SVGs upregulated in adipose tissue and breast glands exhibit minimal gradient values at the cancer boundaries (Supplementary Fig. 10). These observations again suggest that utilizing SVG method alongside StarTrail has the potential to unveil genes linked to previously unidentified domains. It is important to note again that StarTrail is not a SVG method. Complementary to SVG methods, StarTrail highlights the critical role of boundary analysis in spatial omics, revealing its potential to connect omics features with distinct boundary-related changes and uncovering novel aspects of tissue structure, disease development or progression.

### 2 Matching predicted clusters to true layers

We match predicted clusters and true layers by relabeling each predicted cluster to allow easier performance comparison across methods and to enhance result visualization (this method is not involved in the calculation of ARI). When the number of predicted clusters is greater than or equal to the number of true layers, our matching ensures that each true layer has a corresponding set of predicted clusters. In cases where the number of predicted clusters exceeds the number of true layers, some predicted clusters may be merged and assigned to match the same true layer. When the number of predicted clusters matches the number of true layers, a one-to-one correspondence is established.

The matching process starts by iterating through each predicted cluster. Step1: We assign the predicted cluster to a new label based on its best-match (i.e., the highest proportion of overlap) true layer. Step2: As multiple predicted clusters can have the same best-match true layer, we also evaluate the proportion of each predicted cluster in the true layer and select the best-match predicted cluster. For instance, if true layer A is the best-match for both predicted cluster a and b in the first step and true layer B is not assigned to any predicted cluster in the first step, we will examine the proportion of predicted cluster a and b in true layer A. For example, the proportions of predicted clusters a and b in true layer A are 90% and 10%, then true layer A will be assigned to predicted cluster a and true layer B will be assigned to predicted cluster b. Step2 is repeated until all the true layers are assigned to at least one predicted cluster. In order to reduce computation time, we add a threshold *c* for Step2: we only consider predicted cluster when its number of spots is greater than *c*. In our analysis, we used *c* = 10.

**Supplementary Figs. 1.**
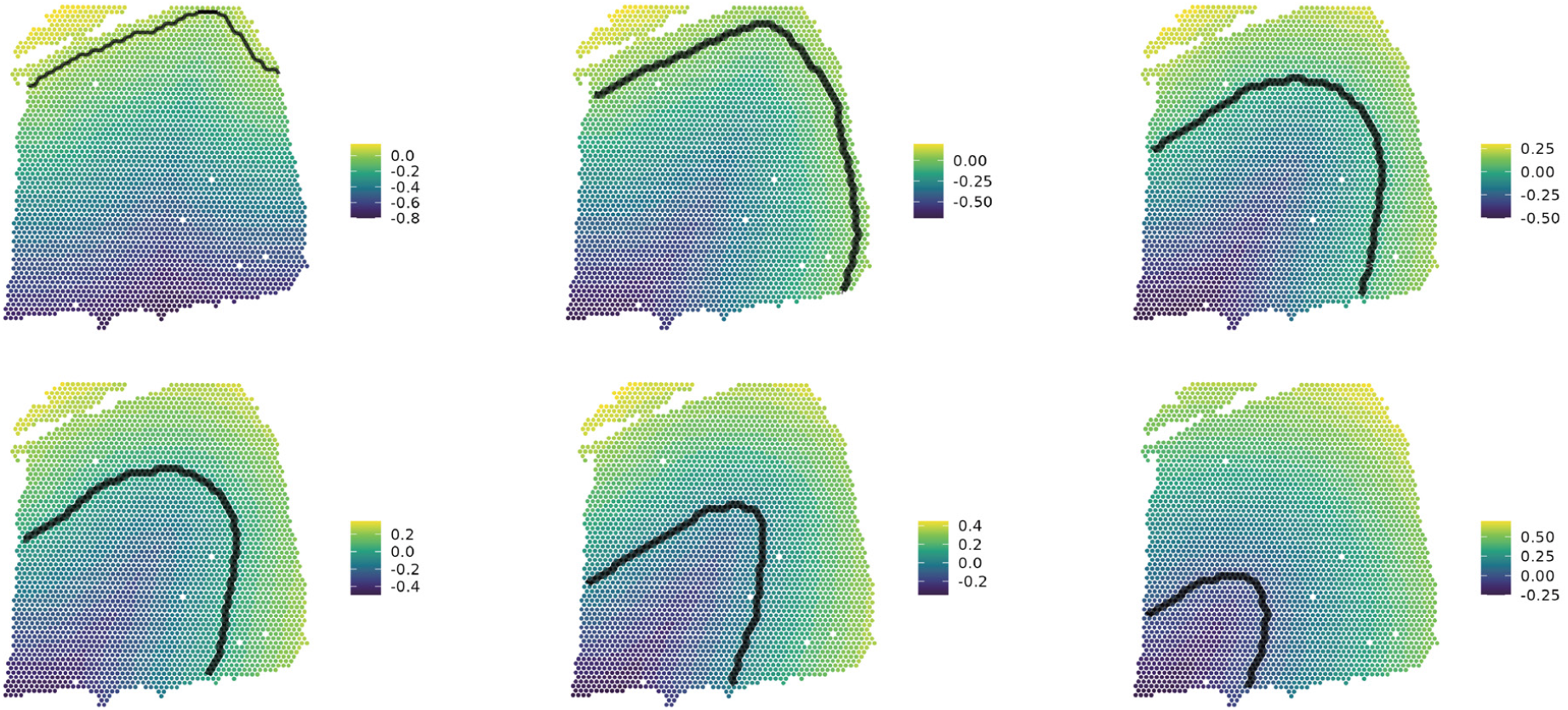
The distance between spots and DLPFC boundary.

**Supplementary Figs. 2.**
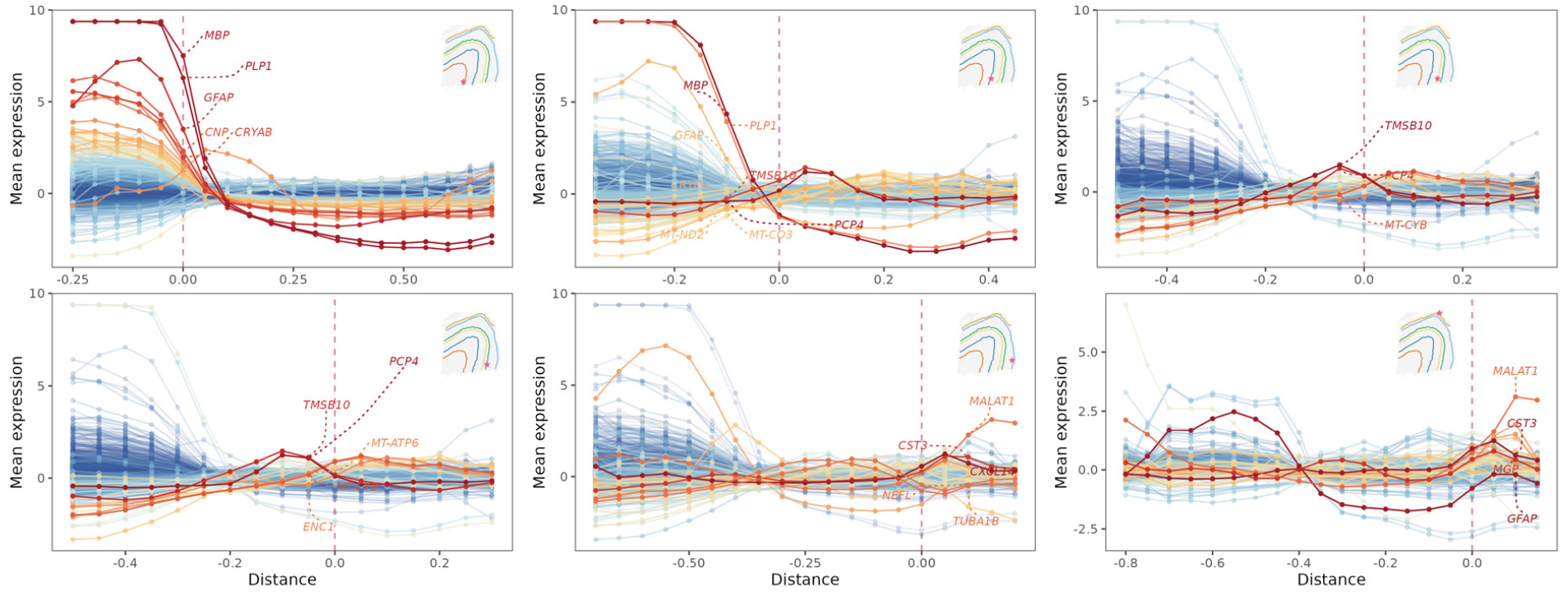
Comparison of mean expression change across each boundary with the gradients in DLPFC analysis. The lines are colored by estimated boundary gradients.

**Supplementary Figs. 3.**
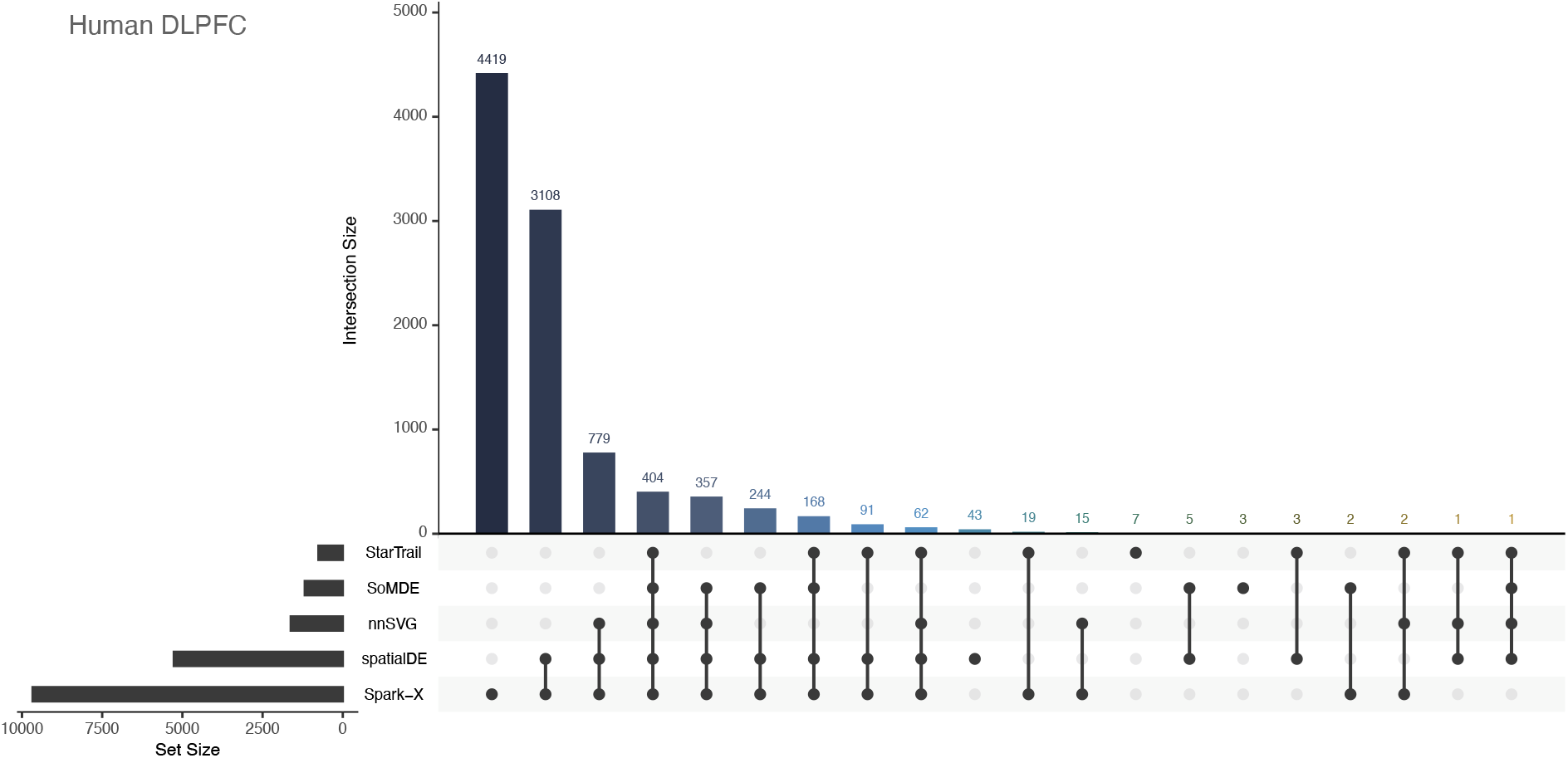
Upset plot for cliff genes and SVGs discovery in DLPFC 151676 analysis. Cliff genes were detected along each of the six boundaries indicated in Fig. 2e.

**Supplementary Figs. 4.**
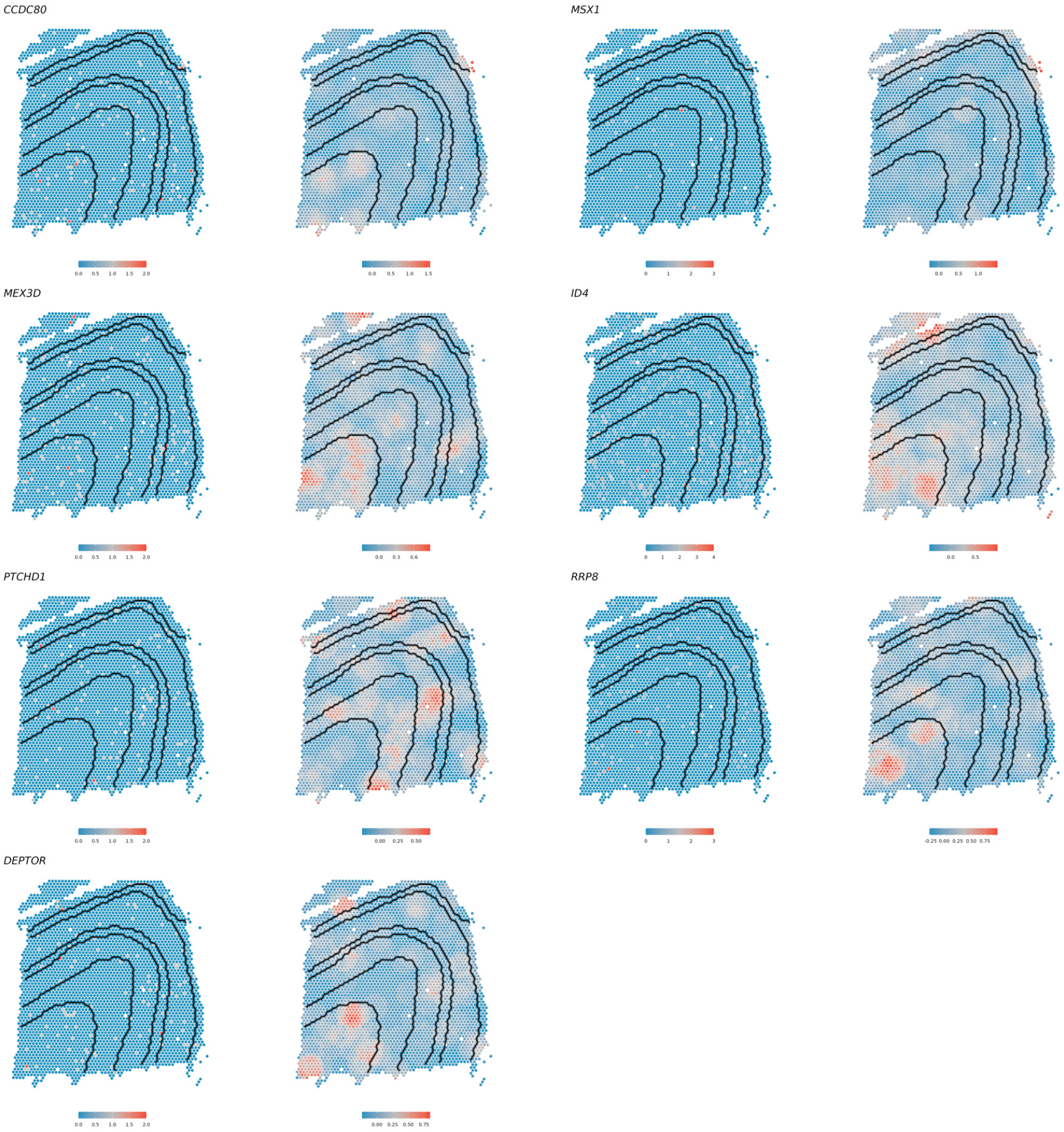
The unique cliff genes detected by StarTrail in DLPFC analysis. For each gene, the left panel is the raw gene expression, the right panel is the smoothed and normalized gene expression.

**Supplementary Figs. 5.**
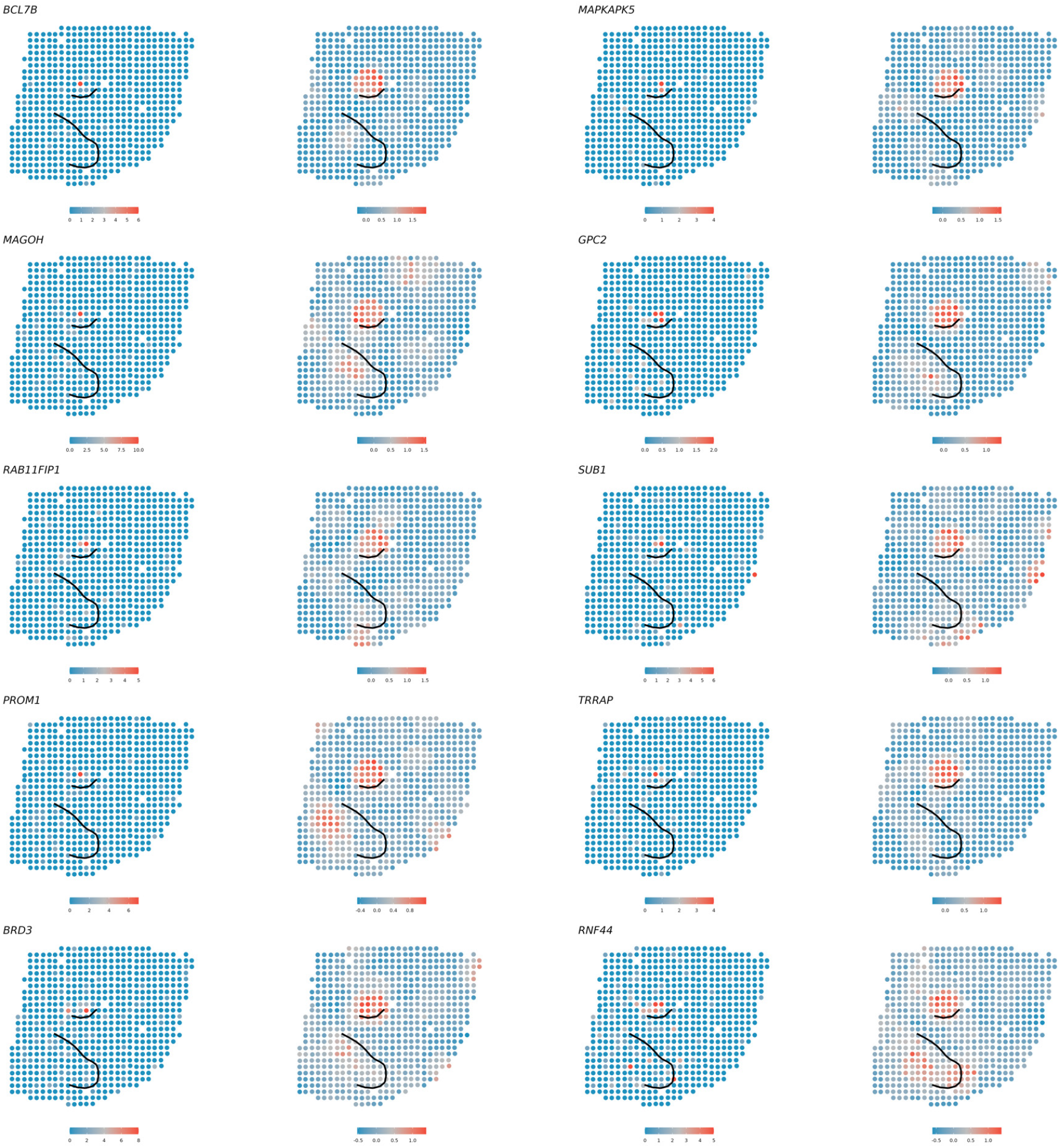
The unique cliff genes detected by StarTrail in HER2+ analysis with top 10 largest absolute gradient on any cancer epithelial boundary. For each gene, the left panel is the raw gene expression, the right panel is the smoothed and normalized gene expression.

**Supplementary Figs. 6.**
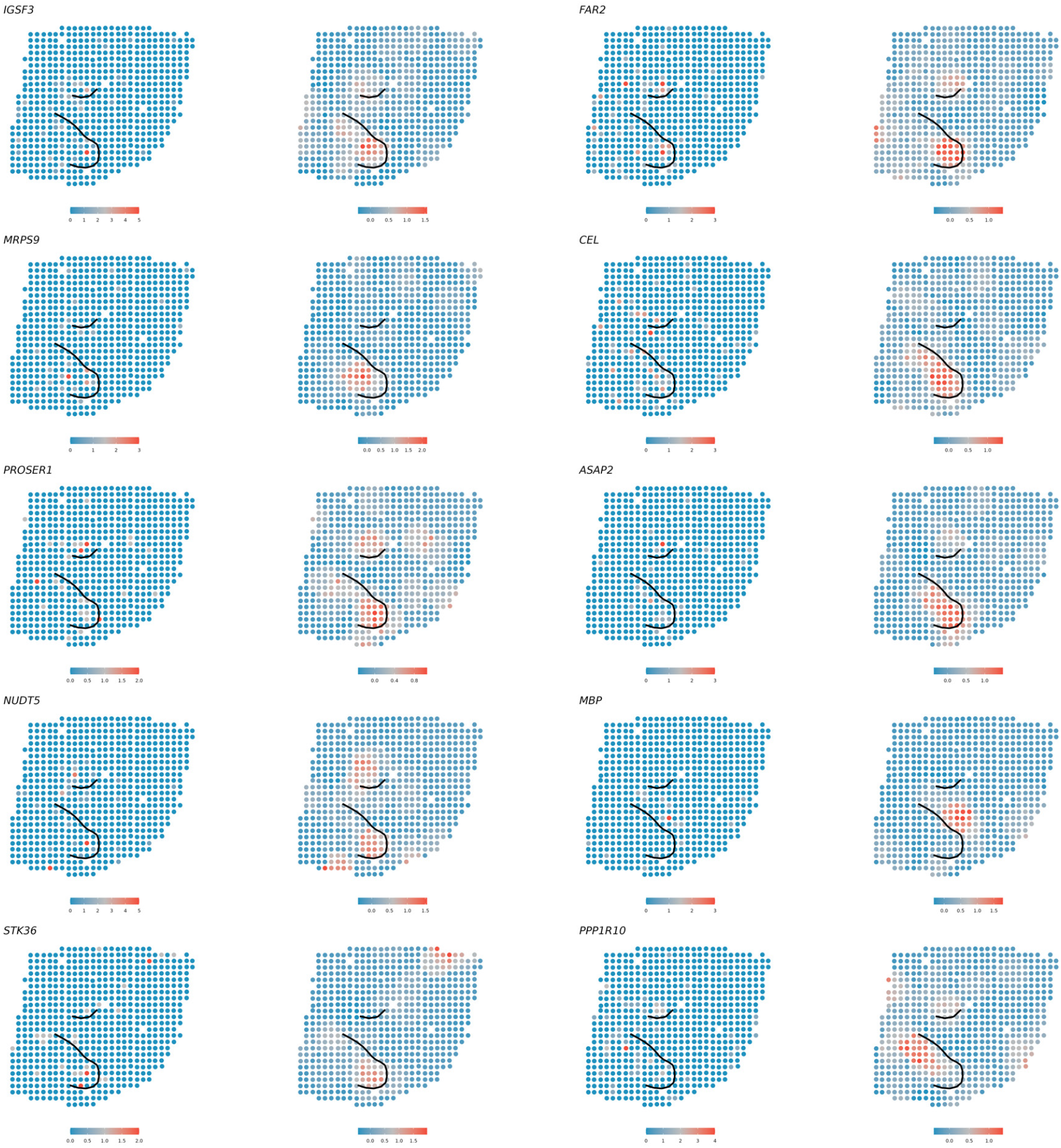
The unique cliff genes detected by StarTrail in HER2+ analysis with top 10 largest absolute gradient on the cancer epithelial multi-boundaries. For each gene, the left panel is the raw gene expression, the right panel is the smoothed and normalized gene expression.

**Supplementary Figs. 7.**
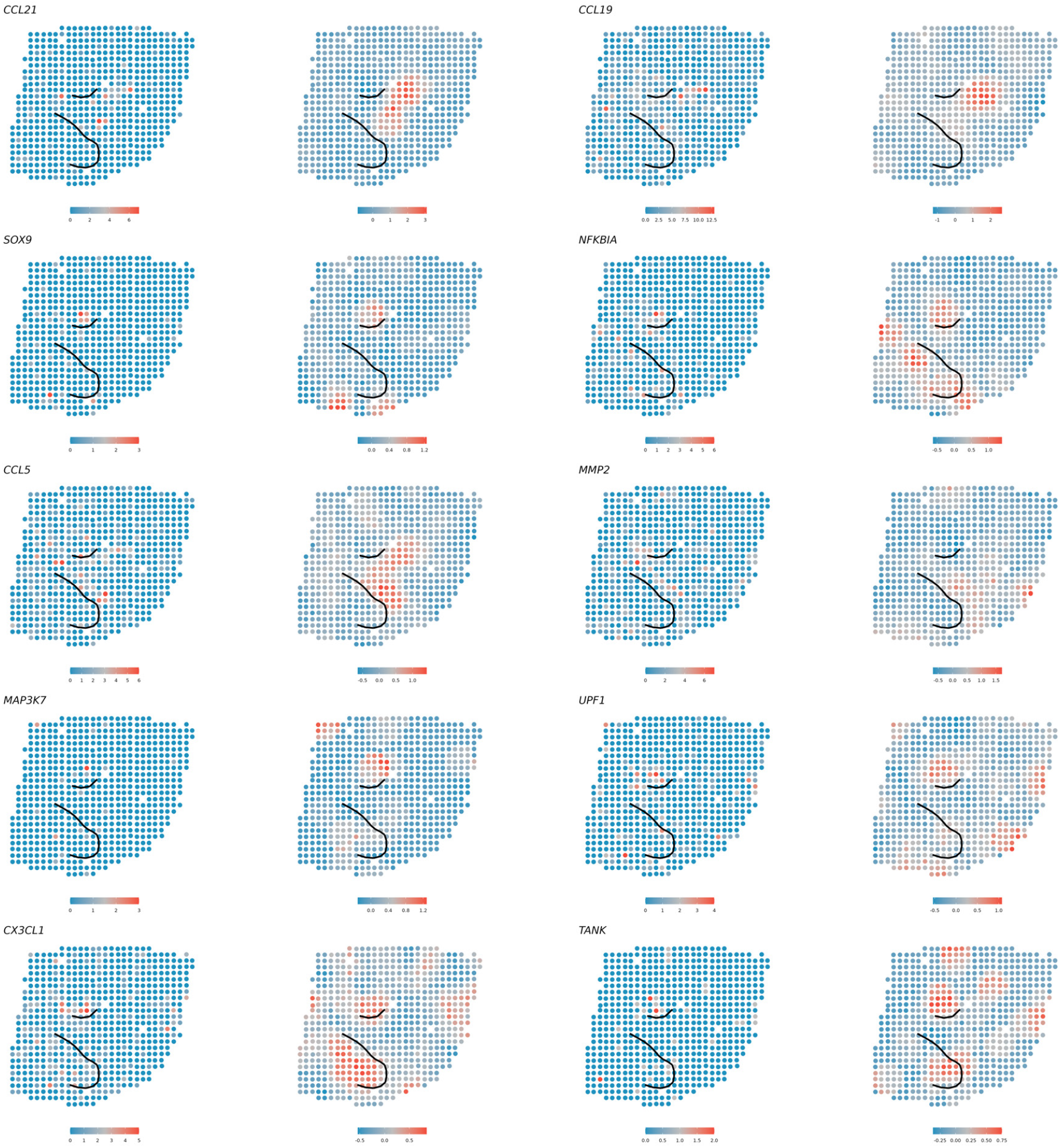
StarTrail detected cliff genes associated with GO:0071347 in HER2+ analysis. For each gene, the left panel shows the raw gene expression, while the right panel shows the smoothed and normalized gene expression.

**Supplementary Figs. 8.**
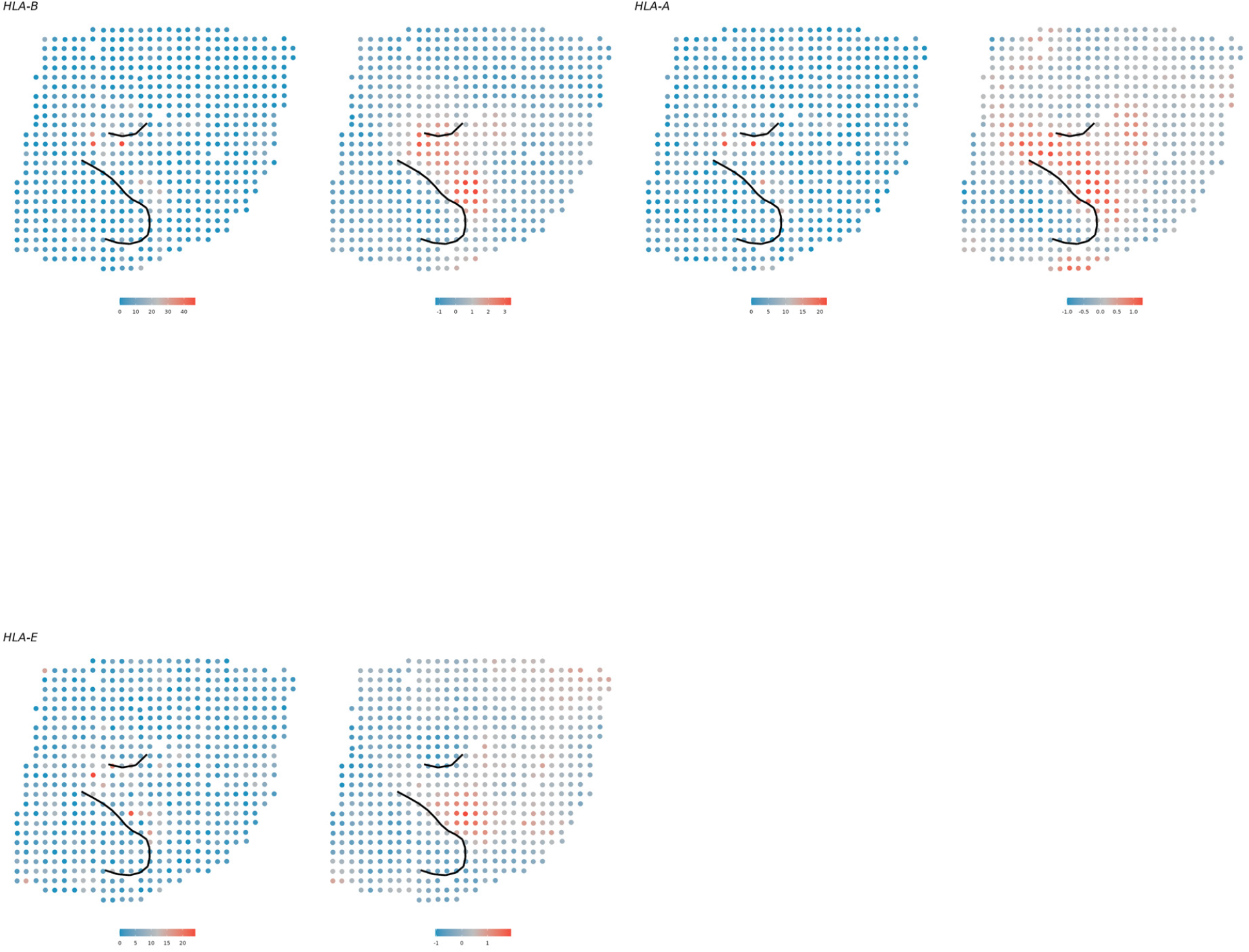
StarTrail detected cliff genes associated with GO:0002715 in HER2+ analysis. For each gene, the left panel shows the raw gene expression, while the right panel shows the smoothed and normalized gene expression.

**Supplementary Figs. 9.**
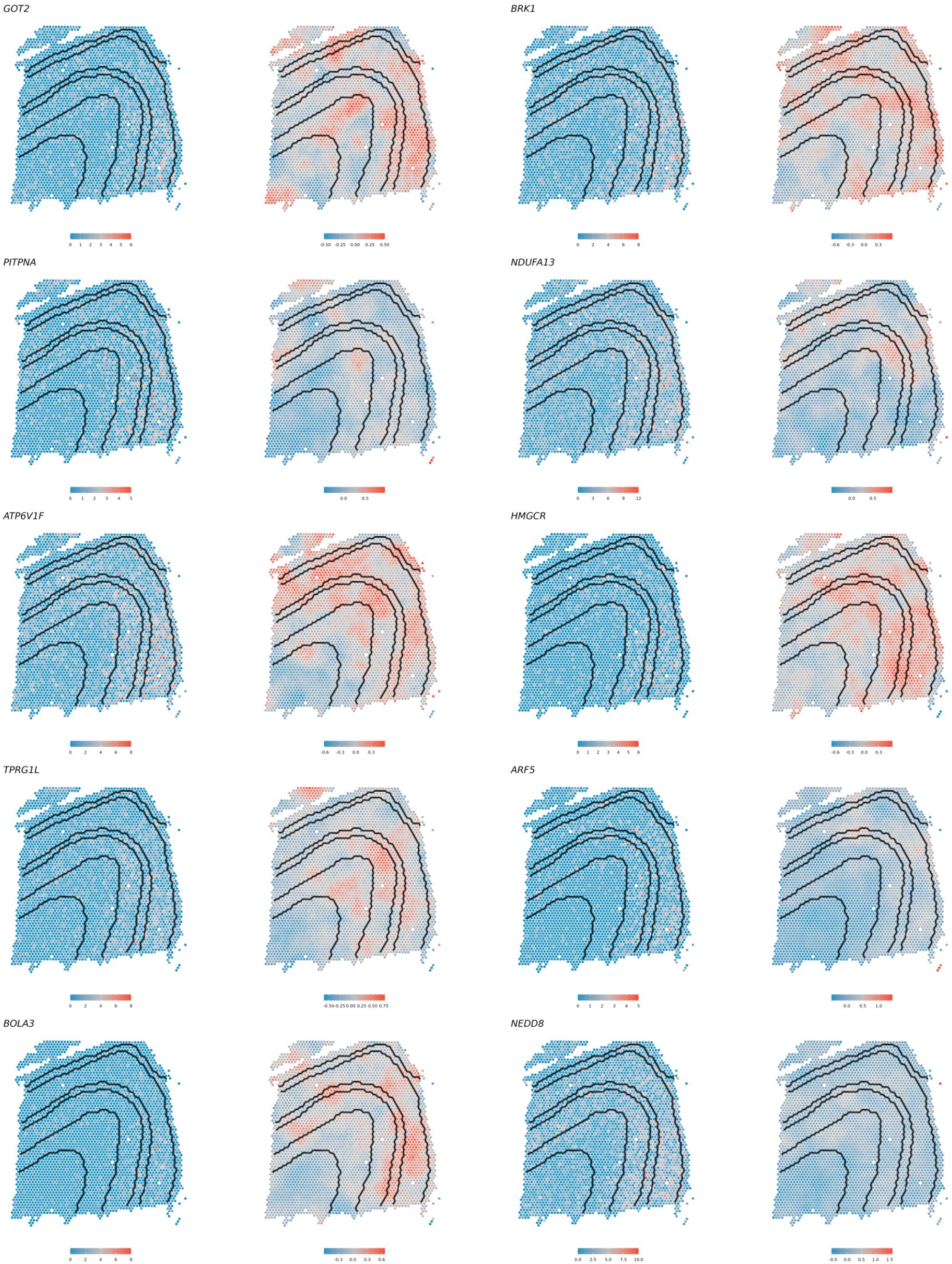
The SVGs genes that are not cliff genes in DLPFC analysis with top 10 smallest absolute gradient on any boundary. For each gene, the left panel is the raw gene expression, the right panel is the smoothed and normalized gene expression.

**Supplementary Figs. 10.**
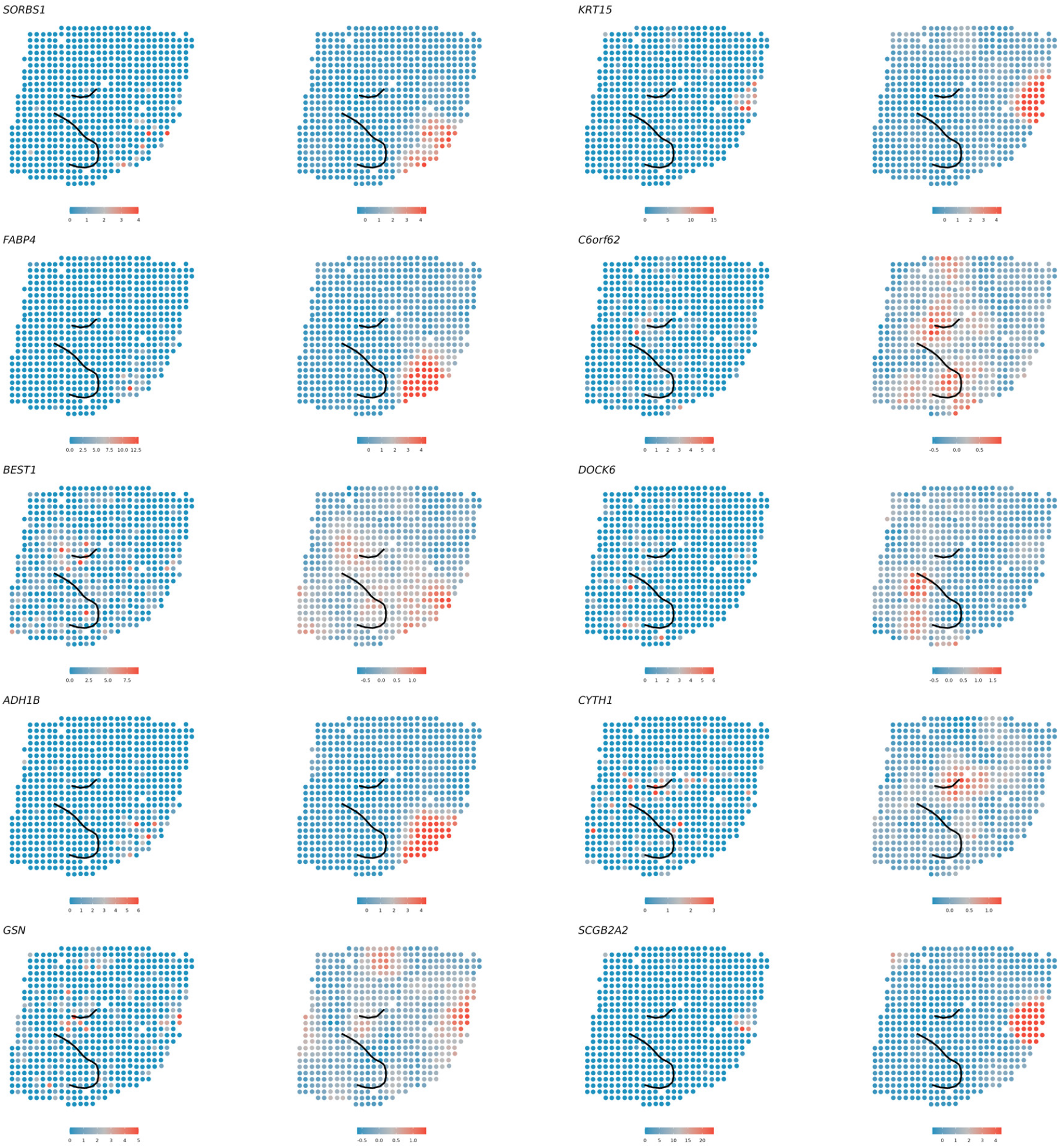
The SVGs genes that are not cliff genes in HER2+ analysis with top 10 smallest absolute gradient on any cancer epithelial boundary. For each gene, the left panel is the raw gene expression, the right panel is the smoothed and normalized gene expression.

**Supplementary Figs. 11.**
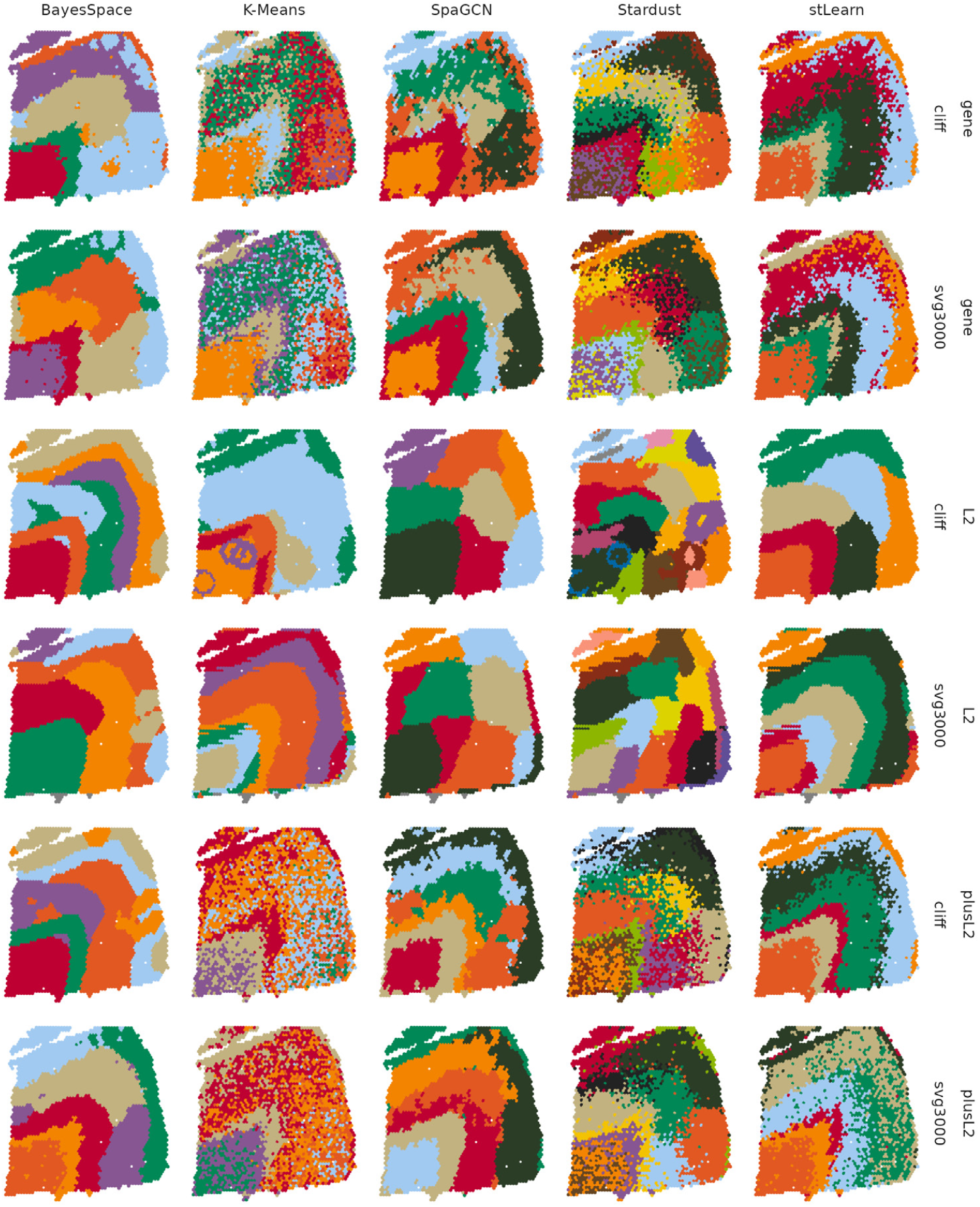
Raw clustering results for all methods with different input (gene expression, *L*^2^, Gene+*L*^2^) in different gene sets (HVG, cliff genes).

## References

[1] Chen, M., Maier, K., Ritenour, D. & Sun, C. Improving global tbi tracking and prevention: An environmental science approach. Global Journal of Health Science 14, 20 (2022).

[2] Zhou, X. et al. Choice of voxel-based morphometry processing pipeline drives variability in the location of neuroanatomical brain markers. Communications Biology 5, 913 (2022).

[3] Lewis, S. M. et al. Spatial omics and multiplexed imaging to explore cancer biology. Nature methods 18, 997–1012 (2021).

[4] Marx, V. Method of the year: spatially resolved transcriptomics. Nature methods 18, 9–14 (2021).

[5] Ståhl, P. L. et al. Visualization and analysis of gene expression in tissue sections by spatial transcriptomics. Science 353, 78–82 (2016).

[6] Stickels, R. R. et al. Highly sensitive spatial transcriptomics at near-cellular resolution with slide-seqv2. Nature biotechnology 39, 313–319 (2021).

[7] Chen, K. H., Boettiger, A. N., Moffitt, J. R., Wang, S. & Zhuang, X. Spatially resolved, highly multiplexed rna profiling in single cells. Science 348, aaa6090 (2015).

[8] Goltsev, Y. et al. Deep profiling of mouse splenic architecture with codex multiplexed imaging. Cell 174, 968–981 (2018).

[9] Sun, S., Zhu, J. & Zhou, X. Statistical analysis of spatial expression patterns for spatially resolved transcriptomic studies. Nature methods 17, 193–200 (2020).

[10] Dries, R. et al. Giotto: a toolbox for integrative analysis and visualization of spatial expression data. Genome biology 22, 1–31 (2021).

[11] Hu, J. et al. Spagcn: Integrating gene expression, spatial location and histology to identify spatial domains and spatially variable genes by graph convolutional network. Nature methods 18, 1342–1351 (2021).

[12] Zhao, E. et al. Spatial transcriptomics at subspot resolution with bayesspace. Nature biotechnology 39, 1375–1384 (2021).

[13] Pham, D. et al. stlearn: integrating spatial location, tissue morphology and gene expression to find cell types, cell-cell interactions and spatial trajectories within undissociated tissues. BioRxiv 2020–05 (2020).

[14] Avesani, S. et al. Stardust: improving spatial transcriptomics data analysis through space-aware modularity optimization-based clustering. GigaScience 11, giac075 (2022).

[15] Cable, D. M. et al. Cell type-specific inference of differential expression in spatial transcriptomics. Nature methods 19, 1076–1087 (2022).

[16] Maynard, K. R. et al. Transcriptome-scale spatial gene expression in the human dorsolateral prefrontal cortex. Nature neuroscience 24, 425–436 (2021).

[17] Li, C., Thijssen, J., Abdelaal, T., Hollt, T. & Lelieveldt, B. Spacewalker: Interactive gradient exploration for spatial transcriptomics data. bioRxiv 2023–03 (2023).

[18] Hildebrandt, F. et al. Spatial transcriptomics to define transcriptional patterns of zonation and structural components in the mouse liver. Nature communications 12, 7046 (2021).

[19] Chitra, U. et al. Mapping the topography of spatial gene expression with interpretable deep learning. bioRxiv (2023).

[20] La Manno, G. et al. Rna velocity of single cells. Nature 560, 494–498 (2018).

[21] Ren, H., Walker, B. L., Cang, Z. & Nie, Q. Identifying multicellular spatiotemporal organization of cells with spaceflow. Nature communications 13, 4076 (2022).

[22] Cang, Z. et al. Screening cell–cell communication in spatial transcriptomics via collective optimal transport. Nature Methods 20, 218–228 (2023).

[23] Hastie, T. & Stuetzle, W. Principal curves. Journal of the American Statistical Association 84, 502–516 (1989).

[24] Banerjee, S. & Gelfand, A. E. Bayesian wombling: Curvilinear gradient assessment under spatial process models. Journal of the American Statistical Association 101, 1487–1501 (2006).

[25] Banerjee, S. & Gelfand, A. E. Bayesian wombling: Curvilinear gradient assessment under spatial process models. Journal of the American Statistical Association 101, 1487–1501 (2006).

[26] Halder, A., Banerjee, S. & Dey, D. K. Bayesian modeling with spatial curvature processes. Journal of the American Statistical Association 1–13 (2023).

[27] Banerjee, S., Gelfand, A. E. & Sirmans, C. Directional rates of change under spatial process models. Journal of the American Statistical Association 98, 946–954 (2003).

[28] Liu, Z. & Li, M. Optimal plug-in gaussian processes for modelling derivatives (2023). 2210.11626.

[29] Datta, A., Banerjee, S., Finley, A. O. & Gelfand, A. E. Hierarchical nearest-neighbor gaussian process models for large geostatistical datasets. Journal of the American Statistical Association 111, 800–812 (2016).

[30] Finley, A. O. et al. Efficient algorithms for bayesian nearest neighbor gaussian processes. Journal of Computational and Graphical Statistics 28, 401–414 (2019).

[31] Wang, F., Bhattacharya, A. & Gelfand, A. E. Process modeling for slope and aspect with application to elevation data maps. Test 27, 749–772 (2018).

[32] Andersson, A. et al. Spatial deconvolution of her2-positive breast cancer delineates tumor-associated cell type interactions. Nature communications 12, 6012 (2021).

[33] Cable, D. M. et al. Robust decomposition of cell type mixtures in spatial transcriptomics. Nature biotechnology 40, 517–526 (2022).

[34] Andersson, A. et al. Single-cell and spatial transcriptomics enables probabilistic inference of cell type topography. Communications biology 3, 565 (2020).

[35] Chen, J. et al. Cell composition inference and identification of layer-specific spatial transcriptional profiles with polaris. Science Advances 9, eadd9818 (2023).

[36] Kennedy-Darling, J. et al. Highly multiplexed tissue imaging using repeated oligonucleotide exchange reaction. European Journal of Immunology 51, 1262–1277 (2021).

[37] Chen, S. et al. Integration of spatial and single-cell data across modalities with weakly linked features. Nature Biotechnology 1–11 (2023).

[38] Caiado, H., Cancela, M. L. & Conceição, N. Assessment of mgp gene expression in cancer and contribution to prognosis. Biochimie (2023).

[39] Jing, X. et al. Cd24 is a potential biomarker for prognosis in human breast carcinoma. Cellular Physiology and Biochemistry 48, 111–119 (2018).

[40] Zhu, J., Sun, S. & Zhou, X. Spark-x: non-parametric modeling enables scalable and robust detection of spatial expression patterns for large spatial transcriptomic studies. Genome biology 22, 1–25 (2021).

[41] Hao, M., Hua, K. & Zhang, X. Somde: a scalable method for identifying spatially variable genes with self-organizing map. Bioinformatics 37, 4392–4398 (2021).

[42] Weber, L. M., Saha, A., Datta, A., Hansen, K. D. & Hicks, S. C. nnsvg for the scalable identification of spatially variable genes using nearest-neighbor gaussian processes. Nature communications 14, 4059 (2023).

[43] Svensson, V., Teichmann, S. A. & Stegle, O. Spatialde: identification of spatially variable genes. Nature methods 15, 343–346 (2018).

[44] Li, Y. et al. Immunoglobulin superfamily genes are novel prognostic biomarkers for breast cancer. Oncotarget 8, 2444 (2017).

[45] Madan, S. et al. Pan-cancer analysis of patient tumor single-cell transcriptomes identifies promising selective and safe chimeric antigen receptor targets in head and neck cancer. Cancers 15, 4885 (2023).

[46] Verschueren, E. et al. The immunoglobulin superfamily receptome defines cancer-relevant networks associated with clinical outcome. Cell 182, 329–344 (2020).

[47] Wu, T. et al. clusterprofiler 4.0: A universal enrichment tool for interpreting omics data. The innovation 2 (2021).

[48] Perrier, S., Caldefie-Chezet, F. & Vasson, M.-P. Il-1 family in breast cancer: potential interplay with leptin and other adipocytokines. FEBS letters 583, 259–265 (2009).

[49] Fix, E. & Hodges, J. L. Discriminatory analysis. nonparametric discrimination: Consistency properties. International Statistical Review/Revue Internationale de Statistique 57, 238–247 (1989).

[50] Cover, T. & Hart, P. Nearest neighbor pattern classification. IEEE transactions on information theory 13, 21–27 (1967).

[51] Rand, W. M. Objective criteria for the evaluation of clustering methods. Journal of the American Statistical association 66, 846–850 (1971).

[52] Hao, Y. et al. Integrated analysis of multimodal single-cell data. Cell 184, 3573–3587 (2021).

[53] Cressie, N. & Wikle, C. Statistics for Spatio-Temporal Data CourseSmart Series (Wiley, 2011). URL https://books.google.com/books?id=-kOC6D0DiNYC.

[54] Banerjee, S., Carlin, B. & Gelfand, A. Hierarchical Modeling and Analysis for Spatial Data Chapman & Hall/CRC Monographs on Statistics & Applied Probability (CRC Press, 2014). URL https://books.google.td/books?id=WVHRBQAAQBAJ.

[55] Stein, M. L. Interpolation of spatial data: some theory for kriging (Springer Science & Business Media, 1999).

[56] Wilmott, P., Howison, S. & Dewynne, J. The mathematics of financial derivatives: a student introduction (Cambridge university press, 1995).

[57] Atkinson, K. An introduction to numerical analysis (John wiley & sons, 1991).

[58] Burden, R. & Faires, J. Numerical Analysis Fourth Edition (PWS KENT Publishing Company Boston, 1989).

[59] Ester, M., Kriegel, H.-P., Sander, J., Xu, X. et al. A density-based algorithm for discovering clusters in large spatial databases with noise, Vol. 96, 226–231 (1996).

[60] Lloyd, S. Least squares quantization in pcm. IEEE transactions on information theory 28, 129–137 (1982).

[61] Blondel, V. D., Guillaume, J.-L., Lambiotte, R. & Lefebvre, E. Fast unfolding of communities in large networks. Journal of statistical mechanics: theory and experiment 2008, P10008 (2008).

[62] Shang, L. & Zhou, X. Spatially aware dimension reduction for spatial transcriptomics. Nature Communications 13, 7203 (2022).

